# Progenitor T cells drive chronic pulmonary type 2 inflammation

**DOI:** 10.1101/2025.10.03.680133

**Authors:** Radomir Kratchmarov, Xiaojiong Jia, Jun Nagai, Hiroaki Hayashi, Kinan Alhallak, Juying Lai, Chunli Feng, Jakob von Moltke, Joshua A. Boyce, Patrick J. Brennan

**Author notes:** These authors contributed equally to this work (Radomir Kratchmarov, Xiaojiong Jia). Corresponding Author: Patrick J. Brennan, M.D., Ph.D. Brigham and Women’s Hospital Hale BTM, 5th floor, Room 5002S 60 Fenwood Road Boston, MA 02115.

## Abstract

Type 2 inflammation unfolds over time and is coordinated by remarkably durable CD4+ Th2 cell responses, standing in contrast to the collapse of adaptive immunity seen in chronic infection and malignancy. In experimental models, short-lived Th2 effector cells and type 2 innate lymphocytes (ILC2) contribute to acute type 2 inflammation, yet the cellular architecture that drives chronic type 2 inflammation and prevents exhaustion remains poorly understood. To define the Th2 landscape in chronic type 2 inflammation, we established a mouse model of long-term pulmonary allergen exposure and found that type 2 inflammation was broadly sustained over at least 4 months despite chronic stimulation. Th2 cells in the lung parenchyma during chronic inflammation were distinct from acute inflammation and resting memory phases, and included an expanded T cell factor-1 (TCF1)-expressing progenitor-like Th2 population. Tissue Th2 progenitors were absent in the nose and in helminth infections, suggesting that type 2 immunopathology is context-specific. Transcriptomic deconstruction of acute and chronic pulmonary Th2 responses defined divergent effector programs as well as the transcriptional signature of a tissue Th2 memory population that resembled human Th2 progenitors. Integrated transcriptomic analysis comparing type 2 inflammation to chronic viral infection revealed a distinct stemness module underlying chronic Th2 inflammation that was separable from a common core memory module. *In vivo*, lung Th2 progenitors coupled self-renewal with effector cell differentiation and could be sustained for weeks without antigen persistence or contribution from the lymph node. Maintenance of Th2 progenitors was partially dependent on lung parenchymal B cells and associated with bronchus-associated lymphoid tissue formation. Moreover, Th2 progenitors, in contrast to Th2 effectors and ILC2s, were sufficient to both initiate and sustain type 2 inflammation. Thus, we define tissue Th2 progenitors as a distinct cellular state that arises during chronic type 2 inflammation and establish the central role of these cells in maintaining the architecture of Th2 responses over time.

## Main Text

Type 2 lymphocytes, including CD4+ Th2 cells and innate lymphoid cells (ILC2), orchestrate type 2 inflammation in barrier tissues during allergic sensitization and helminth infection. These responses are often chronic and involve continuous exposure to antigen, as seen in persistent gastrointestinal parasitic infection and chronic exposure to environmental aeroallergens such as dust and mold. Distinct Th2 populations have been implicated in acute sensitization^1–6^, and in recall responses to episodic exposures by resident memory T cells (TRM) ^7–12^. Th2 TRM secrete cytokines that modulate the local microenvironment, function independently of circulating central memory (TCM),^13^ and mediate resistance to helminth infection through tissue remodeling^14^.

However, it remains unclear how Th2 cells sustain type 2 inflammation during chronic antigen exposure without becoming exhausted. In human allergic diseases with chronic antigen burden, we identified tissue progenitor Th2 cells with self-renewal capacity that mirrors cellular states seen in chronic infection and cancers, and that may be important for long term disease pathogenesis^15^. To functionally test the role of tissue progenitor Th2 cells, we therefore modeled chronic Th2 responses using a mouse model of pulmonary type 2 inflammation that recapitulates key elements of human asthma to better understand the pathways and cell populations sustaining type 2 inflammation (Figure 1a).

**Figure 1:**
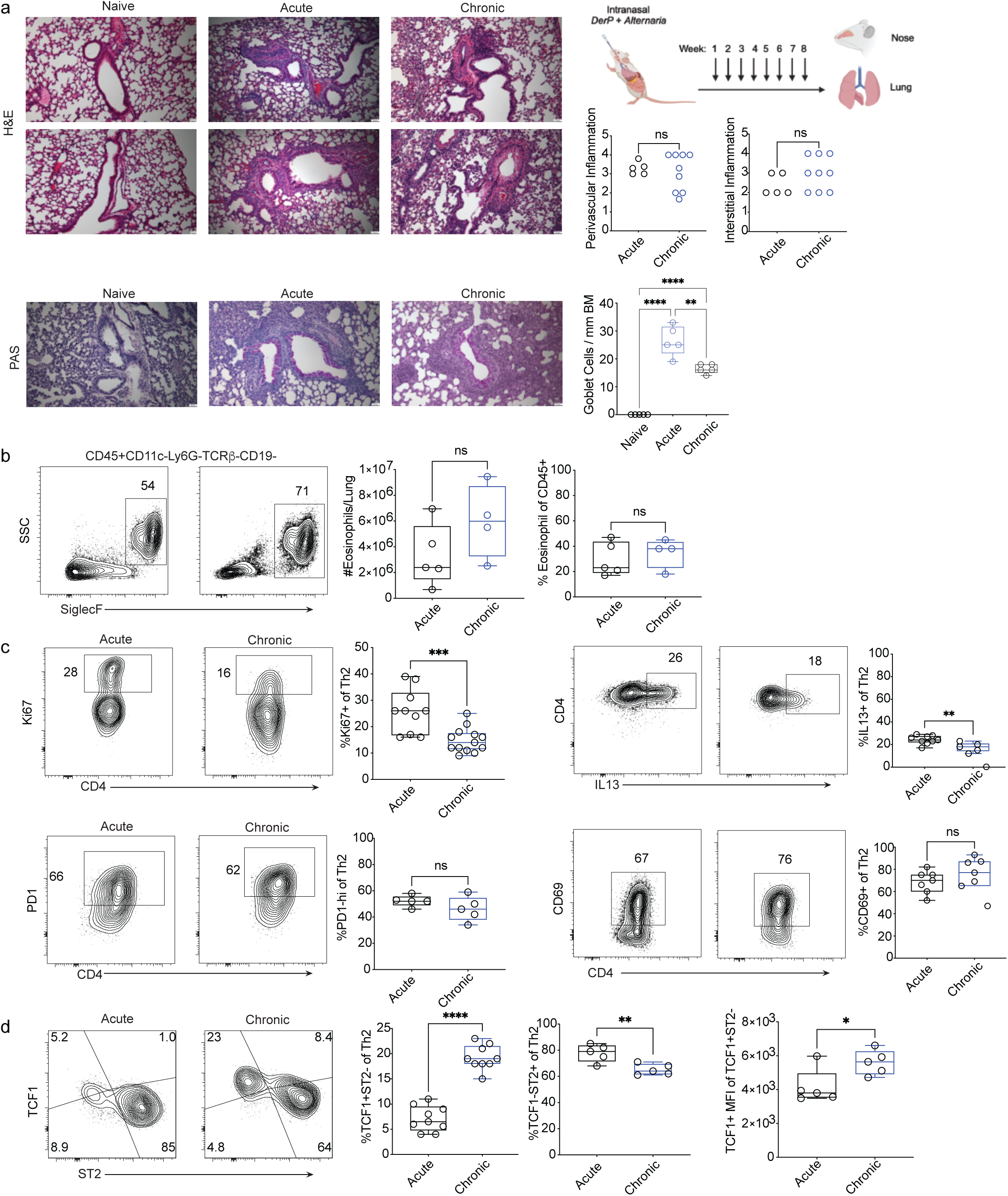
Sustained type 2 inflammation in a model of chronic pulmonary sensitization associated with Th2 heterogeneity. a) Upper: Representative H&E stained images of lungs from naïve, acute (2 weeks) and chronic (8 weeks) time points of C57Bl/6 mice sensitized with *Alternaria*/DerP. Right: quantification of perivascular inflammation and interstitial inflammation through H&E staining at the indicated timepoints. N.s. not significant, One-way ANOVA with Holm-Sidak correction for multiple comparisons. Lower: Representative PAS stained images from naïve, acute sensitization (2 weeks) and chronic sensitization (8 weeks) time points. Right: quantification of numbers of goblet cells per millimeter of basement membrane at the indicated time points. **p<0.01. ****p<0.0001. One way ANOVA with Holm-Sidak correction for multiple comparisons. b) Representative flow cytometry gating of intraparenchymal (CD45 intravascular label negative) eosinophils (CD45+CD11c-Ly6G-Tcrb-CD19-SiglecF+). Middle: Number of eosinophils per lung, Right: percent eosinophils of CD45+ cells per lung, at acute and chromic timepoints. N.s. not significant. Two tailed T test. c) Representative flow cytometry plots of Ki67, IL13, PD1 and CD69 expression at the acute and chronic timepoints for CD4+CD44+CD45(IV)-Foxp3-Gata3+ Th2 cells, with associated quantification of marker positive cells as a percent of CD4+CD44+CD45(IV)-Foxp3-Gata3+ Th2 cells. ** p<0.01, ***p<0.001. n.s. not significant. Two-tailed T test. d) Left: Representative flow cytometry plots of TCF1 and ST2 expression by for CD4+CD44+CD45(IV)-Foxp3-Gata3+ Th2 cells. Middle: Quantification of %TCF1+ST2-progenitor cells of Th2 cells. Right: quantification of %TCF1-ST2+ effector cells of Th2 cells. Far right: quantification of TCF1 mean fluorescence intensity (MFI) among TCF1+ST2-cells. *p<0.05, **p<0.01, ****p<0.0001. Two-tailed T test.

### Sustained chronic pulmonary type 2 inflammation associated with Th2 progenitor differentiation

We performed intranasal sensitization of mice with *Alternaria alternata* (“*Alternaria*”) and house dust mite (*D. pteronyssinus,* “DerP”) extracts weekly for 2 or 8 weeks to model the pathophysiology of human asthma wherein patients are often polysensitized to multiple environmental allergens and encounter distinct stimuli in overlapping fashion. We found robust type 2 inflammatory responses at 2 weeks post-sensitization, as indexed by interstitial and peri-vascular inflammation, as well as goblet cell metaplasia and intraparenchymal eosinophil infiltration (Figure 1a,b). As is evident in human disease, type 2 inflammation was broadly maintained at the 8 week timepoint, with similar levels of cellular infiltration and modestly decreased goblet cell metaplasia. Intravascular labeling distinguished parenchymal T cells and ILC2 from the circulating compartment (Extended Data Figure 1a). Sensitization induced infiltration of both Gata3+ Th2 cells and Gata3+ Tregs^16^. The ILC2 compartment was also comparable between acute and chronic sensitization timepoints, and proliferation levels were similar (Extended Data 1b). To define differences between Th2 cells in the acute and chronic settings, we quantified proliferation (Ki67 expression) and effector cytokine production (IL-13 expression) (Figure 1c). Both proliferation and cytokine production were marginally reduced at the chronic timepoint, but nevertheless persistent. There were no significant differences between expression of the inhibitory receptor and activation marker PD1 or the resident memory / activation marker CD69 at the acute and chronic timepoints. Klrg1, a marker of pulmonary and adipose tissue Tregs with heightened Gata3 expression, was expressed on a subset of Th2s and Tregs (Extended Data Figure 1c), and their frequencies were unchanged over time. Some Th2 cells expressed Klrg1, consistent between the acute and chronic timepoints, and few (<5%) tissue Th2 expressed CD62L at either timepoint (Extended Data Figure 1d). Together, these results indicate that type 2 pulmonary inflammation is broadly sustained over months of sensitization and that Th2 responses exhibit minimal features of exhaustion despite this chronic antigen stimulation. We therefore assessed T cell factor-1 (TCF1) expression to determine the memory/progenitor potential of Th2 cells at different timepoints. We found that TCF1 and ST2 (IL-33 receptor) were reciprocally expressed and marked two broad cellular states: TCF1+ST2-and TCF1-ST2+, as well as a potential transitional TCF1+ST2+ compartment that was present at lower frequency (Figure 1d). TCF1+ST2-were significantly enriched at the chronic timepoint, and moreover, while TCF1+ST2-cells could be found at the acute timepoint, the mean fluorescence intensity of TCF1 was also significantly lower in the acute state. These findings mirror the emergence of TCF1+ stem-like progenitors over time in chronic infection and cancer^17,18^.

We next asked whether differentiation of TCF1+ Th2 cells was context-specific or rather, a broadly conserved feature of type 2 differentiation. We profiled Th2 responses in a commonly used, 4-dose model of acute *Alternaria* sensitization (Extended Data Figure 2a). TCF1 is required for initial Th2 differentiation through repression of interferon-γ and induction of Gata3^19^, and we reasoned that TCF1+ cells might infiltrate the lung early after sensitization. However, in this setting we found that tissue Th2 cells uniformly downregulated TCF1 expression, as compared to bystander Gata3(-) cells and Tregs (Extended Data Figure 2a). Similar results were obtained when we employed a bioinformatic approach by analyzing single cell RNA sequencing data (scRNAseq) from the mouse nose after acute sensitization with *Alternaria*^20^ (Extended Data Figure 2b). We identified clusters of Th2 cells and ILC2 that were expanded following intranasal sensitization, yet Th2 cells uniformly lacked *Tcf7* transcript expression (encoding TCF1) under short-term stimulation (Extended Data 2c and Extended Data Table 1).

We considered other models of chronic type 2 inflammation. The helminth *Heligmosomoides bakeri (H. bakeri)* establishes a chronic infection in the mouse gut, inducing local Th2 responses in the lamina propria and draining mesenteric lymph nodes (MesLN). Mice were infected with *H. bakeri* and the Th2 response was assessed at acute (day 18) and chronic (day 42) timepoints in the gut mucosa and MesLN (Figure 2a,b). In this model, Th2 cells, identified by Gata3 expression, were uniformly CD62L-(Extended Data Figure 2d). Like intestinal ILC2s, Th2 cells in the lamina propria and MesLN expressed the alarmin receptor IL17RB rather than ST2^21^. There was evidence of extensive cell proliferation at both the acute and chronic timepoints (Extended Data Figure 2e). We then assessed TCF1 expression and identified three distinct subsets when stratified by Klrg1 co-expression (TCF1+Klrg1-, TCF1-Klrg1-, and TCF1-Klrg1+). The majority of Th2 in the lamina propria were TCF1(-) at both the acute and chronic timepoints, as compared to the MesLN where TCF1+ cells could be found at both timepoints (Figure 2a). The frequencies of the three major Th2 subsets were unchanged between the acute and chronic timepoints in the mucosa, while TCF1+Klrg1-cells were significantly increased at the chronic timepoint in the MesLN. We observed similar levels of IL17RB expression at both timepoints across the three subsets in both organs. Proliferation was significantly decreased in both the mucosa and MesLN at the chronic timepoint (Figure 2b).

**Figure 2:**
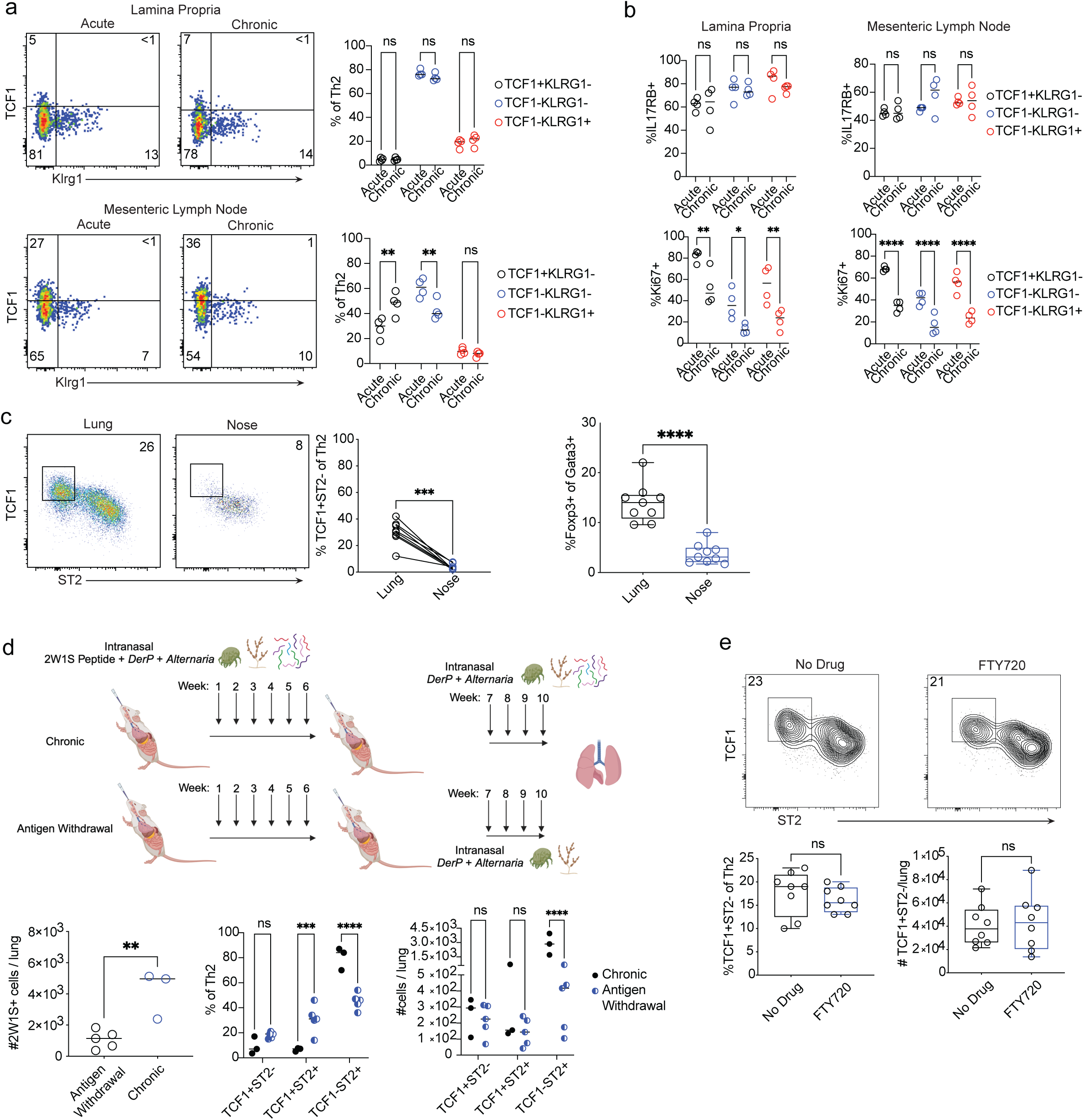
Diverse models reveal site and context specific differentiation of Th2 progenitors. a) Left: expression of TCF1 and Klrg1 by CD4+CD44+Gata3+Foxp3-Th2 cells in the lamina propria (upper) and mesenteric lymph node (lower). Right: quantification of frequencies of TCF1+Klrg1-, TCF1-Klrg1-, and TCF1-Klrg1+ cells. **p<0.01. One way ANOVA with Holm-Sidak’s correction from multiple comparisons. b) Expression of IL17RB+ and Ki67+ cells by each Th2 subset (TCF1+Klrg1-, TCF1-Klrg1-, and TCF1-Klrg1+) in the lamina propria (left) and mesenteric lymph node (right). *p<0.05, **p<0.01, ****p<0.0001. One way ANOVA with Holm-Sidak’s correction from multiple comparisons. c) Left: frequencies of TCF1+ST2-cells in the lung and nose during chronic (8 week) intranasal sensitization. ***p<0.001. Paired T cell test. Right: Percent Foxp3+ Tregs of all Gata3+CD44+CD4+ T cells. ****p<0.0001. T test. Upper: schematic of antigen-specific chronic pulmonary type 2 inflammation model. Lower left: total numbers of 2W1S tetramer positive cells in the lung. following antigen withdrawal or during chronic stimulation. Middle: % indicated subsets of Th2 cells. ***p<0.001, ****p<0.0001. One way ANOVA with Holm-Sidak’s correction from multiple comparisons. Right: number of cells per lung of each Th2 subset. ****p<0.0001. One way ANOVA with Holm-Sidak’s correction from multiple comparisons. e) Left: frequencies of TCF1+ST2-Th2 cells following treatment with no drug or FTY720 treatment. Right: number of TCF1+ST2-Th2 cells per lung. N.s. not significant. T cell test.

These results indicate that chronic helminth infection does not lead to accumulation of a Th2 progenitor population in tissue and suggest that maintenance of long-term responses to these infections may require contributions from secondary lymphoid tissue^22,23^ . We therefore next asked if the tissue microenvironment might influence Th2 progenitor differentiation in our model of chronic pulmonary sensitization. We characterized the Th2 compartment in the nose of mice undergoing chronic DerP/*Alternaria* sensitization in parallel with lung. The nasal Th2 cell phenotype was distinct from that observed in the lung and lacked a substantial TCF1+ST2-progenitor population; Gata3+ Treg infiltration was also lower (Figure 2c). Taken together, these results suggest that tissue Th2 progenitor accumulation is not a broad feature of type 2 responses, but rather that the establishment of this aberrant cell population is context-specific.

We next sought to further investigate the role of TCR signals in Th2 progenitor compartment maintenance by employing a model that allowed for tracking of antigen specific T cell populations using major histocompatibility class II tetramers. We sensitized mice with the peptide 2W1S^16,24^, administered intranasally with DerP/*Alternaria* weekly, and then tracked antigen-specific Th2 cells with fluorescently labeled tetramers (Extended Data Figure 2f). To define the role of TCR signaling in the maintenance of different Th2 subsets, we sensitized mice with 2W1S/DerP/*Alternaria* for 6 weeks and then withdrew 2W1S while continuing DerP/*Alternaria* sensitization for an additional 4 weeks (Figure 2d). After antigen withdrawal, the frequency of TCF1+ST2-cells trended higher though did not meet statistical significance, while the total number of cells remained constant, suggesting that survival/maintenance of Th2 progenitors does not require TCR stimulation. Reciprocally, both the frequency and number of TCF1-ST2+ effector cells decreased after antigen withdrawal. These results indicate that tissue progenitor cells are maintained in the absence of antigen for at least one month at steady state levels, without evidence of contraction. We therefore next assessed whether maintenance of the progenitor subset required ongoing replenishment from a lymph node reservoir. We treated mice with FTY720 to block lymph node egress at the chronic phase of sensitization (Figure 2e). There were no significant differences in the frequency or cell number of TCF1+ST2-Th2 progenitors, despite efficient depletion of T cells from circulation. In settings of chronic inflammation, ectopic lymphoid structures, including bronchus-associated lymphoid tissue (BALT) and tumor-associated tertiary lymphoid structures, can develop^25^ . Indeed, we saw the emergence of lymphoid aggregates consistent with BALT formation during chronic sensitization, which was accompanied by B cell infiltration and germinal center (GC) B cell differentiation (Extended Data Figure 3a-c). As B cells can instruct differentiation and maintenance of tumor-specific stem/progenitor CD8+ T cells^26,27^, we next asked whether B cells could modulate Th2 progenitor cell maintenance. We employed a combination intraperitoneal/direct intranasal protocol for tissue delivery of CD20-directed antibody to facilitate clearance of parenchymal B cells, and found marked reduction of BALT area as well as a decrease in Th2 progenitor cell numbers (Extended Data Figure 3d,e). Together, these results demonstrate that the Th2 tissue progenitor compartment can be uncoupled from antigenic stimulation as well as circulating and lymph node-resident cells for several weeks, and is partially maintained through B cell help in association with the development of ectopic lymphoid structures.

The TCF1+ST2-Th2 compartment arises in the face of chronic antigen stimulation and is associated with ongoing proliferation, cytokine production, and tissue pathology, suggesting that the cellular state of these cells is distinct from that of resting resident memory cells. To directly assess phenotypic differences between putative Th2 progenitors and Th2 resident memory cells, we sensitized mice with DerP/*Alternaria* for 8 weeks and then either continued sensitization for an additional 8 weeks (“Chronic”), withdrew all sensitization for 8 weeks (“Rest”), or withdrew sensitization for 7 weeks and then restimulated with one dose DerP/*Alternaria* (“Recall”) (Extended Data Figure 4a). Tissue inflammation and parenchymal remodeling decreased after two months of rest, while type 2 inflammation was sustained over 4 months of chronic sensitization (Extended Data Figure 4b). Recall responses were brisk, as a single dose of DerP/*Alternaria* was sufficient to re-induce inflammation at levels comparable to chronic stimulation. Tissue eosinophilia resolved during the resting phase and recall and chronic conditions were characterized by similar levels of eosinophilia (Extended Data Figure 4c). We then compared the Th2 compartment across these conditions (Extended Data Figure 4d-f). Frequencies of TCF1+ST2-cells were similar between resting, recall, and chronically stimulated Th2 (Extended Data Figure 4d). We next assessed PD1 expression on Th2 cells, quantifying 3 subpopulations, PD1-hi, PD1-int, and PD1-lo, on both circulating and intraparenchymal cells. Most Th2 were PD1-int or PD1-lo in the resting memory condition, while chronically stimulated and recall Th2 had high levels of PD1 expression suggesting dynamic adaptation to activation (Extended Data Figure 4e). CD69 expression, which marks resident memory T cells but is also a marker of recent TCR stimulation, was also similar between conditions (Extended Data Figure 4f). These surface marker expression patterns assert that the TCF1+ST2-compartment represents a state distinct from resident memory. Taken together these findings across distinct models, organs, and timepoints, indicate that Th2 progenitor differentiation is context-and organ-specific, and may reflect a unique pathway of type 2 lymphocyte responses that is associated with chronic disease.

### scRNAseq/scTCRseq defines the transcriptional signature of Th2 progenitors

While expression of certain key effector molecules and surface receptors differed between acute, chronic, and memory Th2 cells, others were similar, indicating that differences in cell state remained to be defined. To obtain a transcriptome-wide, unbiased profile of Th2 diversity across time, we performed scRNAseq and paired single-cell TCR sequencing (scTCRseq) of activated lung CD4+ T cells at the acute and chronic timepoints after sensitization with DerP/*Alternaria* (Figure 3a, Extended Data Figure 5a), analyzing a total of 17,036 individual transcriptomes after quality control. We identified multiple clusters of Th2 effector cells, which were found in different frequencies at the acute and chronic timepoints (Acute Effector 1/2, and Chronic Effector 1/2) (Figure 3b). The effector clusters expressed high levels of *Il1rl1*, encoding the IL-33 receptor ST2 (Figure 3c). The Acute Effector 2 cluster was predominantly characterized by an interferon response signature, similar to cells identified during acute sensitization with DerP or dog dander^3,28^, while Acute Effector 1 exhibited a transcriptional profile most consistent with classical Th2 effectors, including transcripts for *Il5* and *Il13* as well as *Pparg*^2^. *Il4* expression was relatively increased in chronic effectors (Figure 3c and Extended Data Table 2). Expression of *Il17rb*, encoding the IL25 receptor, and *Nmur1,* encoding a neuropeptide receptor implicated in ILC2 and memory Th2 responses^14,29–31^, were increased in both chronic effector clusters.

**Figure 3:**
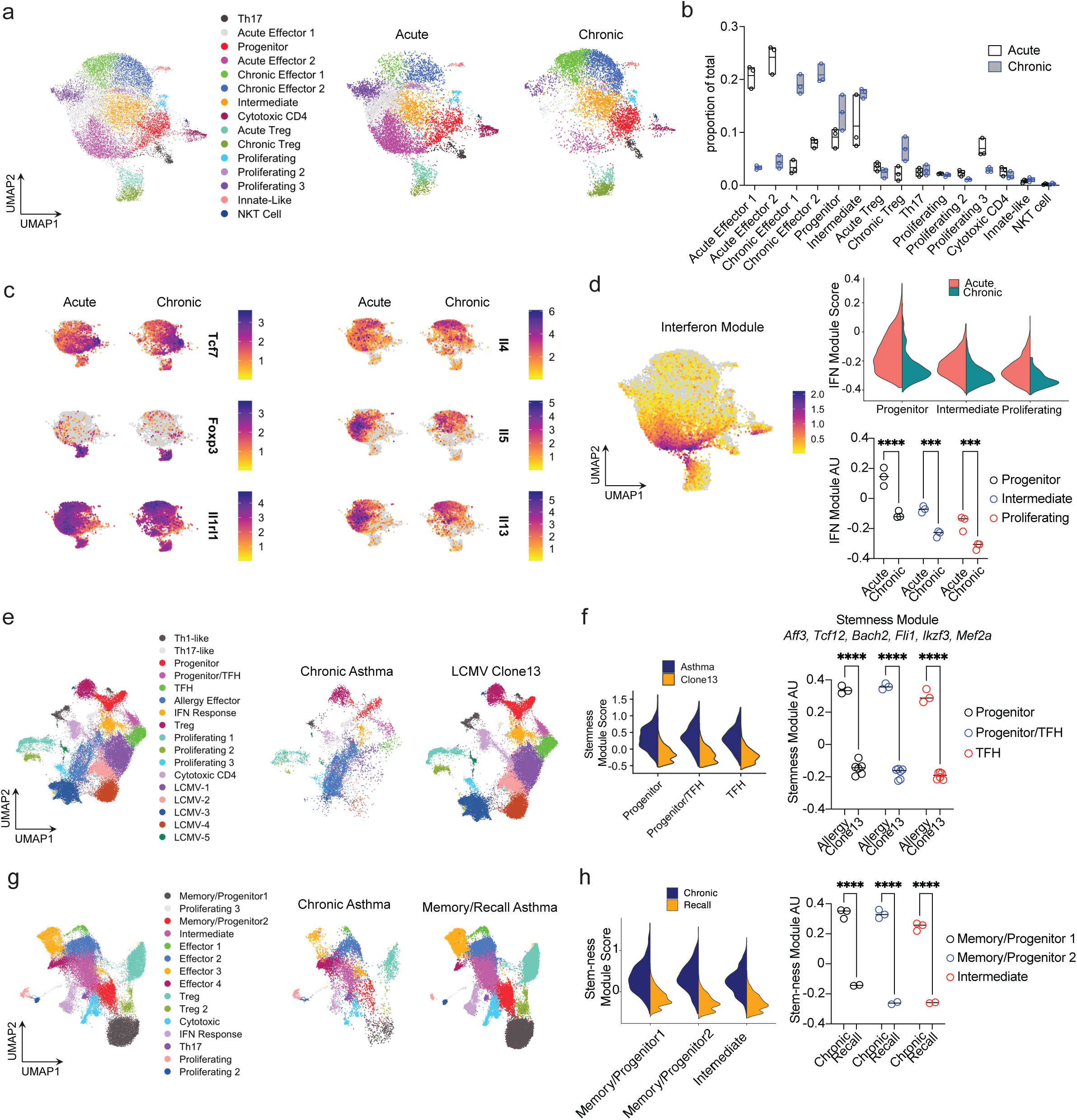
scRNAseq of CD4+ T cells in acute and chronic pulmonary sensitization. a) UMAP of all parenchymal (CD45IV-) lung CD4+CD44+ T cells from acute and chronic timepoint of Alternaria/DerP sensitization. Right: UMAP split by acute and chronic timepoints. b) quantification of frequencies of each cluster at acute and chronic timepoints for each mouse. c) FeaturePlots of key lineage defining transcripts and effector cytokines, split by acute and chronic timepoints. d) Left: FeaturePlot of Interferon module. Upper right: split ViolinPlots of Interferon module for Progenitor, Intermediate, and Proliferating clusters. Lower right: quantification of Interferon module for indicated clusters at acute and chronic timepoints. AU, arbitrary units, ***p<0.001, ****p<0.0001. One-way ANOVA with Holm Sidak’s correction for multiple comparisons. e) Left: UMAP of integrated CD4+ T cells from LCMV Clone13 and Chronic Asthma (week 8 chronic timepoint from Fig3a). Right: UMAP split by disease state. f) Left: Split ViolinPlots of Stemness module for indicated clusters. Quantification of Stemness module for indicated clusters. ****p<0.0001. One-way ANOVA with Holm Sidak’s correction for multiple comparisons. g) Left: UMAP of integrated CD4+ T cells from Resting Memory/Recall Asthma *A. Fumigatus* model and Chronic Asthma (week 8 chronic timepoint from Fig.3a). Right: UMAP split by disease state. Left: Split ViolinPlots of Stemness module for indicated clusters. Quantification of Stemness module for indicated clusters. ****p<0.0001. One-way ANOVA with Holm Sidak’s correction for multiple comparisons.

There were two distinct clusters of Treg cells marked by *Foxp3* expression, one of which was highly enriched at the chronic timepoint (Extended Data Figure 5b). While *Il10* and *Areg* (encoding amphiregulin) expression was largely similar between acute and chronic Tregs, the chronic Treg subset was marked by *Tff1* expression, similar to a recently described Treg state associated with pulmonary fibrosis,^32^ suggesting that chronic Treg responses during type 2 inflammation may be associated with tissue remodeling. There was also broad *Areg* expression by effector Th2 clusters, consistent with prior work suggesting a pathogenic role for amphiregulin in Th2-driven fibrosis^33^. Several clusters containing cells that are not part of the Th2 lineage were also identified, including Th17 cells expressing *Rorc* and *Il23r,* natural killer T (NKT) cells expressing *Zbtb16*, and an innate-like cluster with lower *Cd3e, Cd4, and Lck* expression as well as some *Trdv4* and *Tcrg-C4* expression (Extended Data Figure 5a). A cytotoxic cluster was present and marked by expression of *Eomes* and granzyme transcripts. There were also three clusters defined predominantly by proliferation markers (Proliferating 1-3).

*Tcf7, S1pr1, Slamf6* and *Klf2* expression marked two clusters of less differentiated cells, denoted “Progenitor” and “Intermediate” (Figure 3c and Extended Data Figure 5a). Some cells in the Progenitor cluster expressed *Bcl6*, a transcription factor critical for both T follicular helper and memory/progenitor responses (Extended Data Figure 5b). There was minimal CXCR5 expression at either the acute or chronic timepoints, though *Il21* expression did increase, suggesting that chronic sensitization may be associated with differentiation of T peripheral helper-like cells^34^. Cells in the Intermediate cluster expressed high levels of *Tox*, a transcription factor implicated in exhausted CD8+ T cell responses^35–37^ and moderate levels of *Il1rl1* (Extended Data Figure 5a and Extended Data Table 2).

We then performed pseudobulk analysis to identify differentially expressed transcripts between the main cytokine-producing effector clusters, comparing Acute Effector 1 to Chronic Effector 1 and 2 respectively (Extended Data Figure 5c, Supplementary Tables 3,4). Chronic Effector 1 cells expressed heightened levels of *Il4* as well as the AP1 family transcription factors *Jun* and *Fos,* and Ikaros-family transcripts *Ikzf1* and *Ikzf3*, encoding the transcription factor Ikaros and Aiolos. Acute Effector 1 cells expressed higher levels of *IL5* and *IL13,* as well as the activation marker *Tnfrsf4*, encoding OX40, and inhibitory receptor *Tigit*. Acute Effector 1 cells also expressed higher levels of *Dgat1*, encoding the enzyme diacylglycerol O-acyltransferase 1, implicated in lipid droplet synthesis and ILC2 function^38^. Chronic Effector 2 cells were marked by *Prdm1* and *Ikzf3* expression and had evidence of a differential cytokine production profile including *Il10*.

To define transcriptomic differences that could be associated with self-renewal, we focused on the Progenitor, and Intermediate, and Proliferating 3 (the dominant proliferating) clusters. Progenitor and Intermediate cluster cells during the acute phase expressed higher levels of many transcripts encoding interferon-response elements, including *Ifi27I2a*, *Bst2*, and *Irf7* (Extended Data Figure 6a,b and Supplementary Tables 5-7). There was also higher expression at the acute phase of the CGRP receptor subunit *Ramp3,* recently described to inhibit Th2 differentiation in some contexts^39^. Conversely, *Ramp1* was more highly expressed by Intermediate cluster cells at the chronic phase. Intermediate cells also expressed higher levels of *Il21* after chronic sensitization, which is of note given the role of IL-21 receptor in amplifying Th2 responses in tissue^1^. The folate receptor *Izumo1r*, encoding a folate receptor-like protein^40^ with a role in Treg function^41^,was more highly expressed by Progenitor cells at the chronic phase. Proliferating cluster cells at the chronic timepoint expressed higher levels of *Tcf7* and *Ltb*, in contrast to higher levels of interferon response transcripts at the acute phase (Extended Data Figure 6c). Given the broad trend of elevated interferon signaling pathways during acute sensitization, we constructed an interferon response module based on prior studies of T cells in human inflammatory disease^42^ and scored cells for the Progenitor, Intermediate, and Proliferating 3 (the dominant proliferating cluster) (Figure 3d). Expression of the interferon response module was significantly higher in all three clusters during the acute phase. Together, these results indicate that while conserved clusters can be identified during acute and chronic type 2 pulmonary inflammation, there are divergent cell populations transcriptional patterns that emerge with time.

### Integrated transcriptomic analysis differentiates chronic Th2 progenitors, chronic Th1 progenitors, and Th2 TRM cell states

Progenitor Th2 cells differentiate in the face of chronic antigen burden and share some features with progenitor CD4+ Th1 cells defined in chronic infection^43–45^. During chronic infection, however, CD4+ T cells develop features of exhaustion, which contrasts the ongoing stable type 2 inflammation observed in our model. We therefore undertook a multi-model, comparative bioinformatic approach to identify transcriptomic differences between Th1 and Th2 progenitors by integrating our scRNAseq dataset with two additional datasets examining antigen-specific Th1 cells in the lymphocytic choriomeningitis (LCMV) Clone 13 model (Figure 3e and Supplementary Table 8). In contrast to acute viral infection models, LCMV Clone 13 establishes a chronic infection with ongoing viremia and features of T cell dysfunction, and has been widely used to model the exhausted type 1 inflammatory response. Integration of the datasets revealed an expected divergence in effector cells, with five LCMV-specific effector clusters (LCMV1-5) identified and one “Allergy Effector” cluster (Figure 3e and Extended Data Figure 7a). These clusters had little to no overlap between disease states. Conversely, there was a Treg cluster, an interferon-response cluster, and several proliferating clusters (Proliferating 1-3) that were conserved between the diseases. Moreover, there were 3 identified clusters (Progenitor, Progenitor/TFH, and TFH) identified with significant overlap between the chronic allergen exposure and chronic infection states, and these clusters expressed *Tcf7* and *Slamf6* (Extended Data Figure 7a).

Using our integrated analysis as an anchor point for comparison, we sought to define the transcriptomic differences between the exhausted (Clone13) and chronic allergy (*Alternaria*/DerP) cell states. Many inhibitory receptors and transcription factors have been implicated in various tumor and chronic infection models, and we therefore first assigned a broad “checkpoint” module score to all cells, integrating surface receptor and transcription factor transcripts (see Methods)^46^. The checkpoint module was most highly expressed on LCMV effector cells, as predicted, as well as on Tregs, and generally lowest on progenitors and TFH cells (Extended Data Figure 7b). We then performed pseudobulk analysis to capture differentially expressed transcripts between the exhausted and allergic cell states, focusing specifically on populations with the highest levels of *Tcf7* expression, including the Progenitor and Progenitor/TFH clusters (Extended Data Figure 7c and Supplementary Tables 9,10).

Progenitors from the allergic state expressed higher levels of the glucocorticoid receptor *Nr3c1,* which has been implicated in tissue adaptation and CD8 memory cell fate specification^47^, as well as the NFkB subunit *Nfkb1*. In the allergic setting, the Progenitor cluster also expressed higher levels of *Zeb1*, a transcription factor required for efficient memory cell differentiation^48^, as well as *Stat5b* and *Bach2*, a key regulator of stem-like CD8+ T cells^49,50^. Conversely, Progenitor and Progenitor/TFH cells in chronic infection expressed interferon response genes and the proteasome subunits *Psmb8*/*Psmb9*. Gene set enrichment analysis identified key divergent pathways from the Hallmark collection^51^ (Extended Data Figure 7d). Notably, progenitor cells in chronic asthma showed evidence of IL2/STAT5 signaling, heme metabolism, and UV and androgen responses, while cells from chronic infection redemonstrated elevated interferon responses as well as myc and PI3K/mTOR signaling, further indicative of an anabolic inflammatory state. These results suggest the presence of divergent underlying metabolic programs, of note given the bioenergetic insufficiency seen in chronic infection^52^.

We assessed for differences in key memory transcripts, including *Tcf7* and *Lef1* based on the essential role of these transcription factors in memory/stem-like states, and unexpectedly found no significant differences between chronic infection and chronic type 2 inflammation (Supplementary Tables 9,10). We reasoned that because gene regulatory networks (“regulomes”) defined by intertwined transcription factor modules often govern complex cell states, combinatorial transcription factor sets might better capture divergent cell fate outcomes. Single-cell CRISPR screens have unveiled the roles of such regulomes in controlling CD8+ T cell stemness in cancer^53^, and we therefore constructed a “Core Memory” module and a “Stemness” module based on recent study of tumor-infiltrating CD8+ T cells^54^ (see Methods).

Expression of the Core Memory module was equivalent between the chronic infection and allergic states, suggestive of a core longevity mechanism (Extended Data Figure 7e). However, expression of the Stemness module was significantly enriched in progenitor and progenitor/TFH-like cells in the allergic state (Figure 3f). Conversely, expression of the Interferon response module was also significantly elevated in the setting of chronic infection despite known clearance of interferon complexes (Extended Data Figure 7e)^55,56^, further suggesting a role for interferon signaling in suppressing stemness and promoting exhaustion^43,57^. Taken together, these results highlight a stemness module that can be uncoupled from a shared longevity module between chronic infection and chronic asthma.

Given the transcriptomic divergence between Th2 progenitors and Th1 progenitors in chronic infection, we sought to determine differences between Th2 progenitors during chronic antigen exposure and Th2 cells in the resting memory/recall state. We integrated our scRNAseq dataset with data from an *Aspergillus fumigatus* resting memory/recall model in which type 2 inflammation was induced over several months and in which Th2 resident memory cells were profiled after recall^9^. The integrated bioinformatic analysis identified both conserved and divergent clusters with distinct features (Figure 3g, Extended Data Figure 8a, and Supplementary Table 11). We identified 4 conserved Effector clusters (Effector 1-4), which showed graded expression of the cytokines *Il4, Il5,* and *Il13* (Extended Data Figure 8a). There were two Treg clusters, one of which (Treg 2) was enriched in the memory/recall state. There were also 3 proliferating clusters (Proliferating 1-3), an interferon-response cluster, cytotoxic CD4+ T cells, and Th17 cells, all of which were conserved between the chronic and memory/recall states. Expression of *Tcf7* and *Slamf6* and lower expression of *Il1rl1*delineated two memory/progenitor-like clusters present both during chronic sensitization and memory/recall (Memory/Progenitor 1-2). We applied the same “checkpoint” module upregulated during chronic infection (Extended Data Figure 7b) to chronic and memory/recall type 2 inflammation (Extended Data Figure 8b). Expression of these transcripts was broadly higher in the chronic allergen exposure state, as compared to memory/recall, with the highest levels observed in the effector clusters. These results indicate that chronic sensitization imparts a distinct activation state that distinguishes progenitor cells from tissue resident memory. We then direct compared transcriptional differences between chronic and memory/recall cells in the two memory/progenitor clusters (Extended Data Figure 8c and Supplementary Tables 12,13). Similar to the differences observed between chronic infection and chronic asthma, progenitor cells in both the Memory/Progenitor 1 and Memory/Progenitor 2 clusters expressed higher levels of the glucocorticoid receptor *Nr3c1* as the transcription factors *Aff3, Ikzf3,* and *Tox*.

Conversely, resting cells in the memory/recall state expressed higher levels of the antiapoptotic factors *Bcl2* and *Mcl1*^58^. These patterns suggested possible similarities in the divergent states between chronic infection and chronic asthma, and we therefore applied the Core Memory and Stemness modules, revealing an increase in stemness during chronic allergen exposure compared to recall (Figure 3h and Extended Data Figure 8d). A separate set of transcripts, denoted as the Resting Memory module, was significantly enriched in the memory/recall state (Extended Data Figure 8d). Thus, progenitor Th2 cells that differentiate during chronic sensitization are transcriptionally divergent from Th2 resident memory cells and are defined by a stemness module which distinguishes them from cell states seen in chronic infection and resting memory.

### scTCRseq highlights Th2 clonal lineages and putative differentiation patterns

Integrated scRNAseq analysis suggested that progenitor cells enriched during chronic sensitization could serve as a cellular reservoir for the replenishment of Th2 effectors. To address these putative developmental relationships, we performed scTCRseq in parallel with scRNAseq transcriptomic analysis to establish clonal relationships. TCR sequences were used as genetic barcodes to track cells across clonal expansion^15^. Large and hyperexpanded clones were detected at both the acute and chronic timepoint and could be seen across clusters (Extended Data Figure 9a). Correspondingly, acute and chronic effectors had the least clonal diversity (Extended Data Figure 9b). We then assessed clonal overlap between clusters in the Th2 lineage, omitting innate/invariant cells and Th17 cells (Extended Data Figure 9c). The largest degree of clonal overlap was observed among the Effector (Acute and Chronic) clusters as well as with the Intermediate cluster. There was also substantial overlap between the Progenitor cluster and each of the Effector and Intermediate clusters. Clonal overlap between Th2 effectors, Cytotoxic cells, and Tregs was limited, as was overlap between Progenitors and Tregs. We then calculated a clonal expansion index between clones common to the Progenitor cluster and each other cluster to infer clonal expansion directionality^15^ (Extended Data Figure 9d). This index is based on the concept that a T cell clone proliferates as it terminally differentiates into effector cells, such that there should be more cells of the differentiated state present, and positive expansion indices are consistent with the starting node occupying a less differentiated state. Using the Progenitor cluster as the candidate starting node, we observed statistically significant positive clonal expansion indices into the 4 effector clusters as well as the Intermediate cluster. Visualization of individual TCR clones in the UMAP space further supported the notion of the Progenitor cluster as a less differentiated cell state, with clonal relations to Th2 effector cells (Extended Data Figure 9e). Clonal expansion was neutral for overlapping Progenitor/Treg clones as well as Progenitor/Cytotoxic clones (Extended Data Figure 9d).

### TCF1+ST2-Th2 progenitors self-renew, populate the effector compartment, and sustain pulmonary disease

As phenotypic, transcriptomic, and TCR clonal analyses suggested that the lung parenchyma harbored Th2 cells with progenitor potential, we next used an adoptive transfer approach to directly test this hypothesis. After 8 weeks of chronic stimulation with DerP/*Alternaria*, candidate progenitor (Ly108+ST2-), intermediate (Ly108+ST2+), and effector (Ly108-ST2+) Th2 populations were sorted and transferred into naïve congenic hosts (Figure 4a). Ly108 was used as a cell surface surrogate of Tcf1 expression for these experiments (Extended Data Figure 10a)^59,60^. Following adoptive transfer, mice were sensitized for 2 weeks with DerP/*Alternaria,* and the fate of the engrafted cells was analyzed (Figure 4b). All 3 transferred populations were readily identified 2 weeks after transfer and sensitization, with significantly higher numbers of progenitor-derived cells present in the lungs and mLN compared with transferred effector cells. Corresponding to the observed cell numbers, Ki67 staining identified increased proliferation in the transferred progenitor Th2 as compared to the intermediate and effector populations (Extended Data Figure 10b). The fate of each transferred population differed, with progenitor Th2 cells yielding progenitor, intermediate, and effector phenotype cells, while transferred effector cells only produced further effector Th2 cells. Transferred intermediate cells differentiated into intermediate and effector populations (Figure 4b). These data suggested that the identified progenitor population indeed has functional progenitor potential, coupling self-renewal with effector generation. These results also support a model in which differentiation from progenitor to effector is unidirectional.

**Figure 4:**
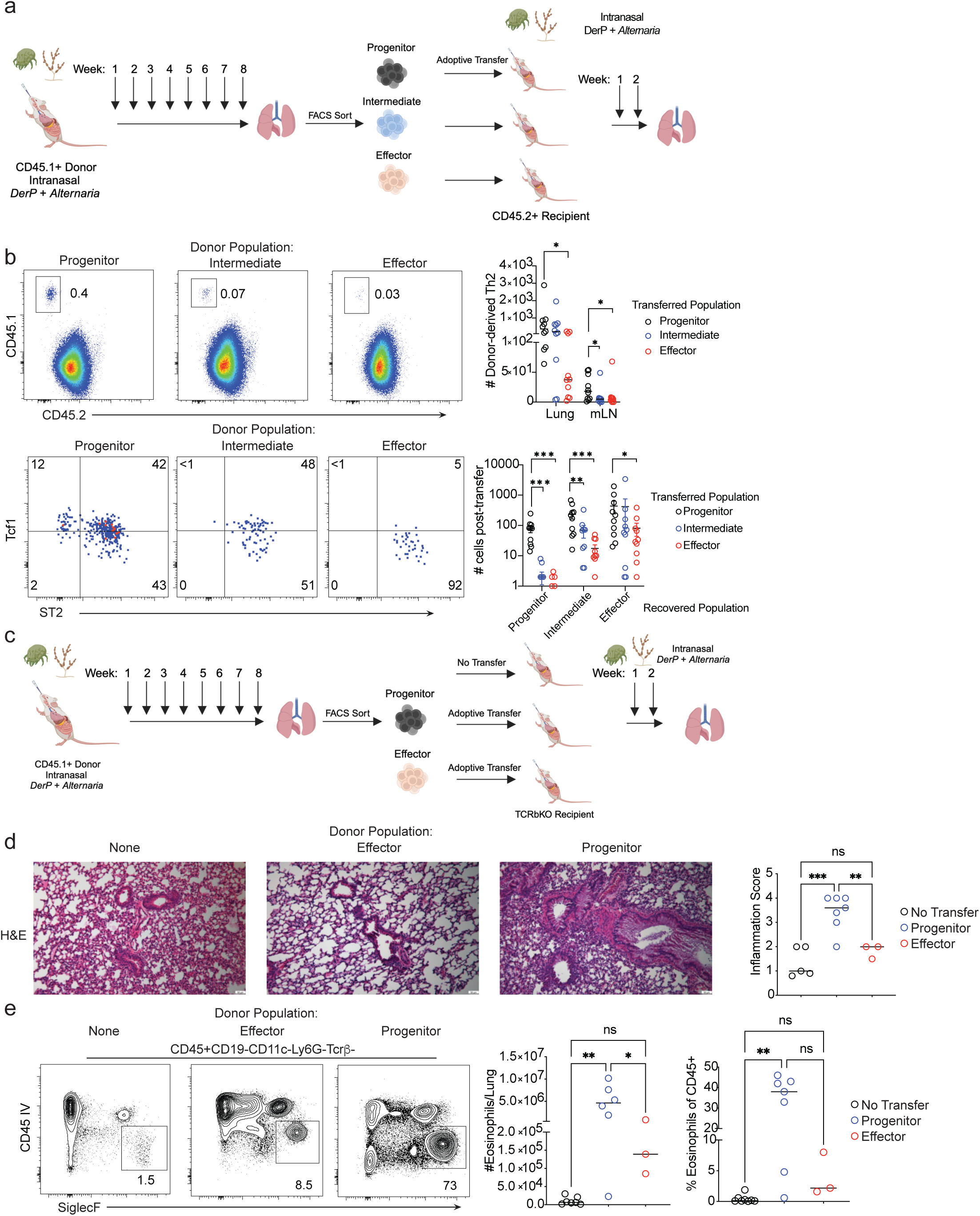
Th2 progenitors self-renew and differentiate into effector cells to initiate and sustain type 2 pulmonary inflammation. a) Schematic of adoptive transfer model of cell fate. b) Upper Left: representative flow cytometry gating of transferred CD45.1+ donor Progenitor, Intermediate, or Effector cells, distinguished from CD45.2+ host cells. Upper Right: quantification of total cell numbers recovered from each donor population in the lung and mediastinal lymph node (MLN). *p<0.05, all others not significant. One-way ANOVA with Holm-Sidak’s correction for multiple comparisons. Lower Left: representative flow cytometry gating of TCF1 and ST2 expression by Gata3+Foxp3-CD45.1+ donor cells. Lower Right: Quantification of numbers of recovered cells of each phenotype following transfer of donor cells of each phenotype. c) Schematic adoptive transfer model of disease causality with TCRbKO recipient mice. d) Representative H&E stains of lungs following adoptive transfer of the indicated populations. Right: quantification of inflammation score. ***p<0.001, n.s. not significant. One-way ANOVA with Holm-Sidak’s correction for multiple comparisons. e) Left: representative flow cytometry gating of parenchymal eosinophils following adoptive transfer of indicated Th2 populations into TCRbKO recipient mice. Middle: quantification of total number of eosinophils per lung and Right: quantification of eosinophils as a percent of CD45+ cells in the lung. *p<0.05, **p<0.01, n.s. not significant.

To assess the functional contribution of the progenitor Th2 population, we next asked whether progenitor cells were sufficient to initiate and sustain lung inflammation. We sorted progenitor (Ly108+ST2-) or effector (Ly108-ST2+) Th2 populations from mice sensitized with DerP/*Alternaria* for 8 weeks, transferred these cells to TCRβ-deficient mice that lack ab T cells yet still have ILC2 and B cells, and sensitized recipient mice for 2 additional weeks (Figure 4c). Mice that received progenitor cells showed robust type 2 inflammation including goblet cell hyperplasia and interstitial alveolar infiltrate (Figure 4d). Only transferred progenitor cells led to ILC2 cell proliferation, a readout of cross-cellular activation thought to contribute to type 2 inflammation in the lung^24^ (Extended Data Figure 10c). Transfer of progenitor cells also led to prominent eosinophilic infiltration that was significantly higher than that induced by transferred Th2 effector cells (Figure 4e). These data demonstrate that Th2 progenitors are sufficient to confer robust type 2 inflammation, while Th2 effectors induced only limited tissue eosinophilia.

It is becoming increasingly appreciated that aberrant stem/progenitor CD4+ T cell responses are critical to the pathogenesis of autoimmune^61–65^ and type 2 inflammatory disease^15^. Reciprocally, stem/progenitor CD4+ cells are required to sustain the response to chronic infection^60^. These mirror-image cell states constitute key therapeutic targets for disease-modifying cellular therapies. Although the factors that sustain these states remain unclear, further studies will clarify the interplay between Th2 cells and the tissue microenvironment, including specialized DC subsets^66^, cytokine-rich niches^23,67^, and cross-talk with ILC2^68,69^, all of which may modulate chronic inflammation. Our data identify chronic type 2 inflammation as a driver of progenitor-like Th2 cells exhibiting transcriptomic features of stemness, layered onto a core memory module, and with the capacity to sustain type 2 lung inflammation in the face of ongoing antigen exposure. Thus, our data now outline a functionally essential progenitor Th2 cell state that initiates and sustains type 2 inflammation and whose transcriptional signature can be uncoupled from conventional memory and exhausted T cells.

## Methods

### Mice, Chemicals, and Allergic Sensitization

Mouse strains in this study include B6 CD45.1 (B6.SJL-*Ptprc^a^ Pepc^b^*/BoyJ), TCRbKO (B6.129P2-*Tcrb^tm1Mom^*/J), and wild type B6, all from the Jackson Laboratory. *Alternaria alternata* extract and *Dermatophagoides pteronyssinus* extract were purchased from Stallergenes Greer. 6ug of DerP and 12.5ug *Alternaria* were used for each sensitization. Mice were anesthetized with isofluorane, and extracts were applied after dilution in normal saline. Adoptive transfers and intravascular antibody labeling were all performed via retroorbital injection. 10,000-20,000 cells of the indicated Th2 populations were adoptive transferred to each recipient mouse. 2W1S peptide (amino acid sequence: EAWGALANWAVDSA) was obtained from GenScript (Piscataway, NJ, USA) and reconstituted at 2mg/mL in PBS. Trace amounts of NaOH were added dropwise to the PBS to facilitate solubility of 2W1S in a slightly alkaline environment. For experiments with FTY720 (Sigma), the compound was dissolved in 7.5% DMSO, 92.5% H2O at 3mg/mL to generate stock solutions and then diluted further in H2O prior to intraperitoneal dilution for a final dose of 20ug/mouse (1mg/kg). FTY720 was administered every 48 hours for two weeks. For B cell depletion experiments, mice were treated with anti-CD20 antibody (clone MB20-11, BioXCell) at 250ug I.P for one dose, followed by 50ug intranasal doses every 48 hours for 2 weeks.

### Lung Histology

Lung lobes were rinsed in PBS and then fixed for 24 hours in 4% paraformaldehyde and embedded in paraffin. Sections were stained by hematoxylin and eosin (H&E) to quantify inflammatory cell infiltrate. A blinded pathologist imaged bronchovascular bundles from at least 6 different fields for each lung, and cellular infiltration was graded on a scale of 0-4, with 0 denoting no inflammation and 4 indicating severe inflammation. Bronchus-associated lymphoid tissue area was also quantified as square millimeters of lung parenchyma, in blinded fashion. To evaluate goblet cell metaplasia, periodic acid-Schiff (PAS) staining was performed on 5mm paraffin sections. Goblet cells were quantified in the bronchial epithelium of at least four independent bronchovascular bundles for each lung. Data are presented as goblet cells per mm of basement membrane.

### Nasal mucosa preparation

Mouse snouts were harvested from euthanized mice by initially making incisions along both sides of the lower jaw, followed by a circumferential cut at the skull base to detach the head. The skin was carefully removed, and the skull was isolated from the surrounding muscle and brain tissues. The zygomatic arches were excised, and longitudinal incisions along the frontal bone exposed the nasal cavity. Under a dissecting microscope, nasal mucosa was scraped away using a scalpel until only bone remained visible. The collected mucosa was then incubated at 37 °C for 60 mins in 750 µl of 10% FBS-RPMI medium containing 2 mg/ml collagenase type IV (Worthington) and 10µg/ml DNase I (Sigma). After incubation, the digested mucosa was vortexed for 30 seconds and further dissociated by triturating with an 18-gauge syringe needle, followed by a 21-gauge needle. The cell suspension was passed through a 70µm cell strainer, washed once with FACS buffer, and prepared for extracellular staining.

### Helminth Infection and Tissue Preparation

*Heligmosomoides bakeri* (*H.bakeri*) larvae were raised and maintained as previously described^70^. For *H.bakeri* infections, mice were infected by oral gavage with 200 L3 larvae and euthanized at acute (day 18) or chronic (day 42) timepoints to collect tissues for analysis by flow cytometry. For small intestine lamina propria preparations, similar methods to Mayer *et al.* were used^71^. Briefly, mice were anesthetized with 2,2,2-Tribromoethanol (Avertin) (Sigma) and the small intestine was nicked at the duodenum and transected at the cecum and flushed with 20 mL 37°C HBSS (no Ca2+/ Mg2+) + 10mM HEPES. The mice were then perfused through the heart with 30mL of 30mM EDTA + 10mM HEPES in HBSS (no Ca2+/Mg2+). Three minutes after initiating perfusion, the first 10 cm of the proximal SI were harvested, Peyer’s patches were removed, tissue was fileted open, cut into 2-3 cm sections, and transferred to 35mL ice cold HBSS (no Ca+2/Mg+2) + 10mM HEPES and shaken vigorously for 30 seconds to release epithelial cells. Intestinal pieces were filtered through a mesh and stored in HBSS + 5% FCS on ice and then transferred into pre-warmed digest buffer composed of RPMI (with Ca2+/Mg2+) supplemented with 20% fetal calf serum (FCS) (Biowest), 1mg/ml collagenase A (Sigma Aldrich), 10 mM HEPES, 1 mg/ml DNase I (Sigma Aldrich) and shaken at 200 rpm at 37°C for 30 minutes. Tissues were vortexed and cells were passed through a 100µm filter, washed with ice cold HBSS (no Ca2+/Mg2+) + 10mM HEPES, then passed through a 40µm filter, and washed with HBSS (no Ca2+/Mg2+) + 10mM HEPES. The cells were then centrifuged for 5 minutes at 1500 rpm, and supernatant discarded. Pelleted cells were then washed and stained for flow cytometry. For preparation of mesenteric lymph nodes (mesLN), lymph nodes were resected, stored in HBSS + 5% FBS on ice, mashed through a 70 µm filter into dPBS and washed once with dPBS.

### Flow cytometry

For intracellular staining of transcription factors and cytokines, cells were fixed/permeabilized for 25 minutes at room temperature with the FoxP3 Transcription Factor Staining Buffer kit (eBioscience). All intracellular staining was performed at 4°C for 1 hour. For assessment of responses to cytokine stimulation, cells were stimulated with PMA/ionomycin for 4 hours. All samples were run on a BD Fortessa or BD Symphony. Data was analyzed using FlowJo (v10).

### Single-cell RNA and TCR sequencing

Cells from mouse lungs were isolated after sensitization with Alternaria and DerP at the indicated times and sorted on a BD FACSAria Fusion using a 70μm nozzle. Single-cell suspensions were loaded at a cell density of 50,000 cells per chip using the Chromium Single-Cell Next GEM 5’ Reagent Kit (V3, 10x Genomics) according to the manufacturer’s instructions. Gene expression matrices were generated using Cell Ranger. We removed low quality cells with <200 measured genes and a high percentage of mitochondrial transcripts (>12%). All subsequent steps were performed using standard functions in Seurat (NormalizeData, FindVariableFeatures, ScaleData, RunPCA, FindNeighbors, FindClusters, RunUMAP) with default parameters and resolution 0.5. The analysis package scCustomize was used for generation of some plots^72^.

Single cell TCR sequencing (scTCRseq) analysis, TCR sequences were associated with each cell barcode and analyzed using standard functions in the package scRepertoire^73^. Clonal overlap was visualized as a Circos plot using the package Circlize. Individual TCR clones/cells were visualized on the UMAP space in black highlight. To calculate the pairwise expansion indices for overlapping clones, we divided the number of cells expressing a TCR sequence in the putative effector cluster by the number of cells expressing that TCR sequence in the root cluster. Positive values indicate clonal expansion in the effector cluster, whie negative values indicate clonal expansion in the root cluster.

### Integrated bioinformatic analysis

Several publicly available scRNAseq datasets were analyzed including: acute Alternaria sensitization (GSE245074)^20^, resting memory/recall *Aspergillus fumigatus* sensitization (GSE190795)^9^, and LCMV Clone13 infection (GSE181474^60^ and GSE182320^44^). For all analyses of previously published datasets, we download the available CellRanger pipeline files and then analyzed via the same Seurat pipeline as for our scRNAseq studies. We removed low quality cells with <200 measured genes and a high percentage of mitochondrial transcripts (>12%). All subsequent steps were performed using standard functions in Seurat (NormalizeData, FindVariableFeatures, ScaleData, RunPCA, FindNeighbors, FindClusters, RunUMAP) with default parameters and resolution 0.75. For comparisons of chronic type 2 inflammation and chronic infection or resting Th2 memory, respectively, we integrated the scRNAseq datasets using the Harmony^74^ algorithm to mitigate batch effect with parameters 11 = 0.3 and θ = 4. Transcripts included in the respective modules included:

- Core Memory module: *Tcf7, Lef1, Foxo1, Sell, S1pr1, Satb1, Ets1, Pou2f2*
- Stemness module: *Aff3, Tcf12, Bach2, Fli1, Ikzf3, Mef2a*
- Resting memory module: *Bcl2, S1pr1, Mcl1, Klf2, Ly6a*
- Interferon response module: *Ifi27l2a, Ifi27, Ifitm1, Ifitm2, Ifi203, Ifitm1, Isg15, Ifit3, fit2, Ifit1, Isg20, Iigp1, Irf7, Bst2, Oas1a, Oas2, Oas3, Trim56, Trim22, Rnf213*

### Statistical analysis

Statistical analyses were performed using Prism (GraphPad). Individual statistical tests are indicated in each figure legend as well as significance cutoffs. All error bars shown are representative of the standard error (SEM) unless otherwise noted. All statistical tests for hypothesis testing were two-sided.

### Data Availability

The newly generated scRNAseq data will be deposited at the GEO publicly repository and available for public access on publication. All analysis and figures were generated using publicly available software packages, without the need for any custom code.

## Acknowledgements

We thank Nicole A. Case and Hadas T. Pahima (Brigham & Women’s Hospital) for assistance with retroorbital injections, as well as to Junning Case and the BWH Center for Cellular Profiling Single Cell Genomics Core for assistance with single-cell transcriptomics.

---

**Extended Data Figure 1:**
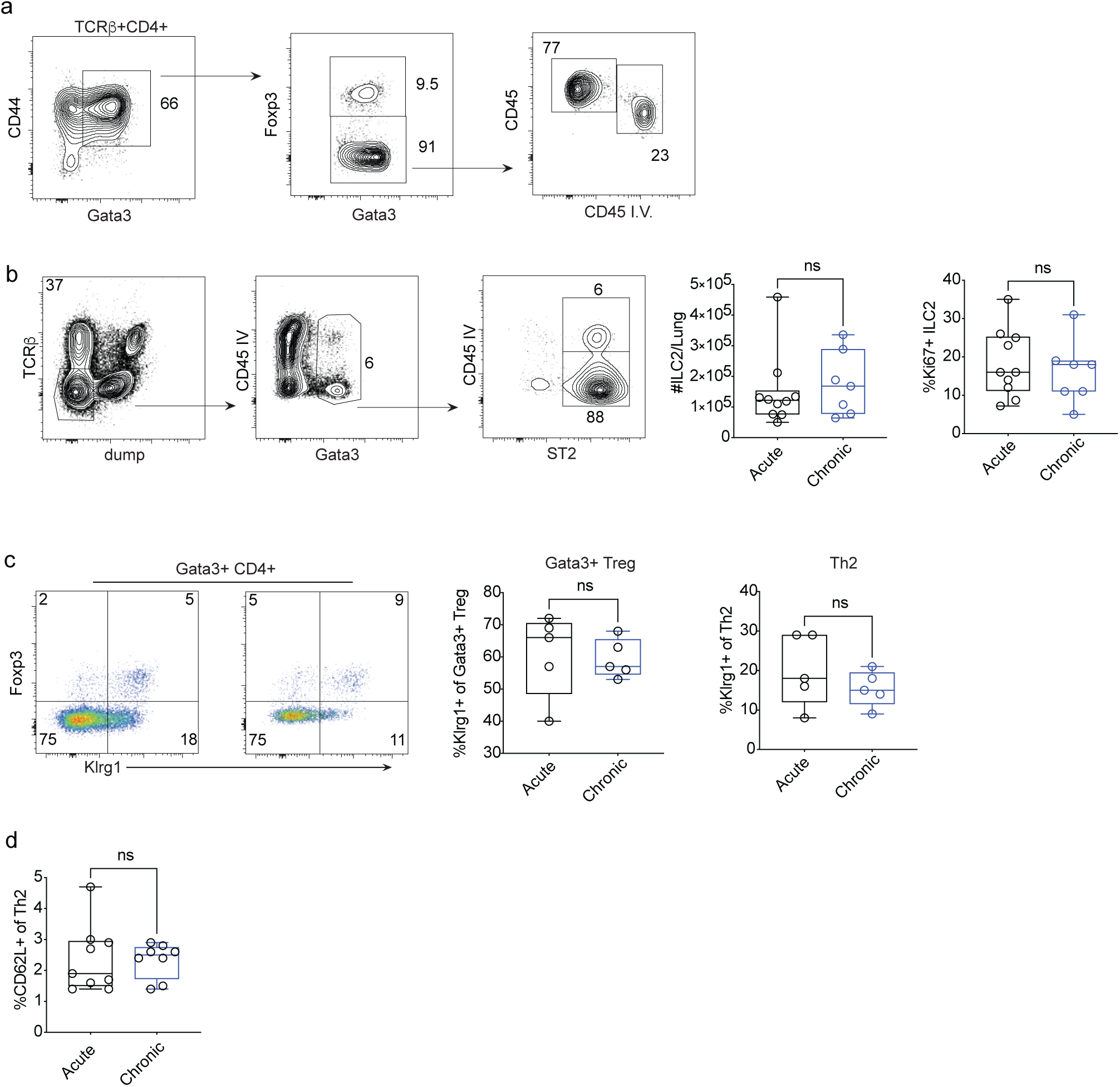
A chronic model of pulmonary allergen sensitization. a) Representative flow cytometry gating sequence for identification of intraparenchymal Gata3+ Th2 cells. b) Left: Representative flow cytometry gating sequence for identification of intraparenchymal ILC2 cells. Middle: quantification of total number of ILC2s in the lung at acute and chronic sensitization timepoints. Right: quantification of proliferating ILC2 frequencies as a perfect of total ILC2s in the lung at indicated timepoints. N.s. not significant. Two-tailed T test. c) Left: Representative flow cytometry gating of Klrg1 expression by Foxp3+Gata3+ Tregs and Foxp3-Gata3+ Th2 cells. Middle: %Klrg1+ cells of Gata3+ Tregs. Right: %Klrg1+ cells of Th2 at the acute and chronic timepoints. N.s. not significant. Two-tailed t test. d) quantification of CD62L+ cells as a percent of Th2 at the acute and chronic timepoints. N.s. not significant. Two-tailed t test.

**Extended Data Figure 2:**
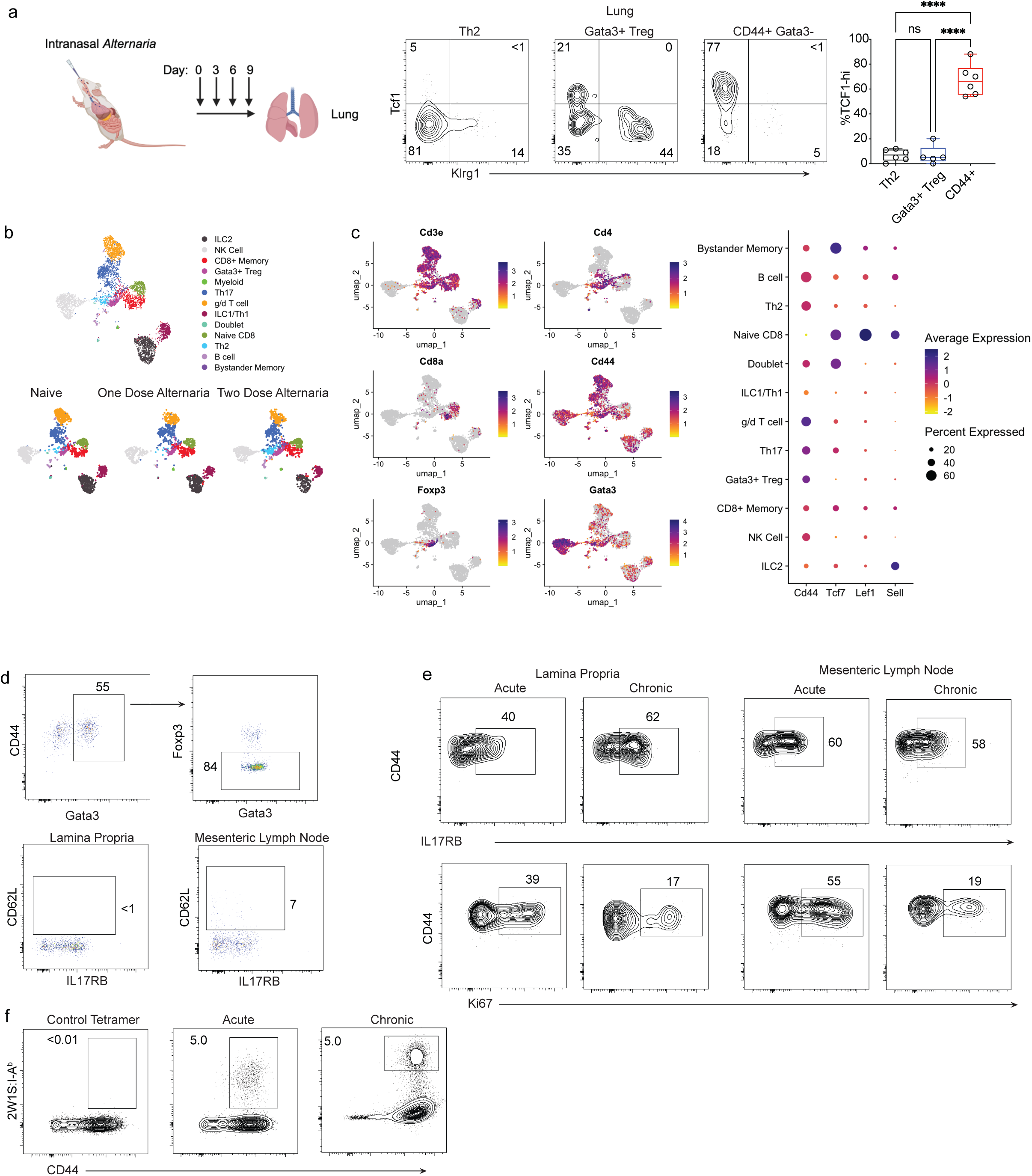
Distinct models of type 2 inflammation. a) Left: Schematic of acute *Alternaria* sensitization model. Middle: Representative gating of TCF1+ cells among Th2 cells (CD4+CD44+CD45(IV)-Foxp3-Gata3+), Gata3+ Tregs (CD4+CD44+CD45(IV)-Foxp3+Gata3+), and other memory effector CD4+ T cells (CD4+CD44+Gata3-). ****p<0.0001. n.s. not significant. One way ANOVA with Holm-Sidak’s correction for multiple comparisons. b.) UMAP of lymphocytes from the Ualiyeva *et al.*^20^ scRNAseq dataset. Right: UMAP split by treatment condition (naïve, one dose Alternaria, two dose Alternaria). c) Left: FeaturePlots of key lineage defining transcripts. Right: DotPlot of key progenitor and effector markers d) Upper: Representative flow cytometry gating of Th2 cells in the lamina propria and mesenteric lymph nodes of mice infected with *H. bakeri*. Lower: Expression of CD62L and IL17RB by Th2 cells in the indicated organs. e) Representative flow cytometry gating of IL17RB (upper) and Ki67 (lower) expression by Th2 at the acute and chronic timepoints in the lamina propria and mesenteric lymph nodes. f) Representative 2W1S tetramer staining in the lung for CD4+CD44+CD45(IV)-Foxp3-Gata3+ Th2 cells at the indicated timepoints after sensitization with DerP/*Alternaria*/2W1S.

**Extended Data Figure 3:**
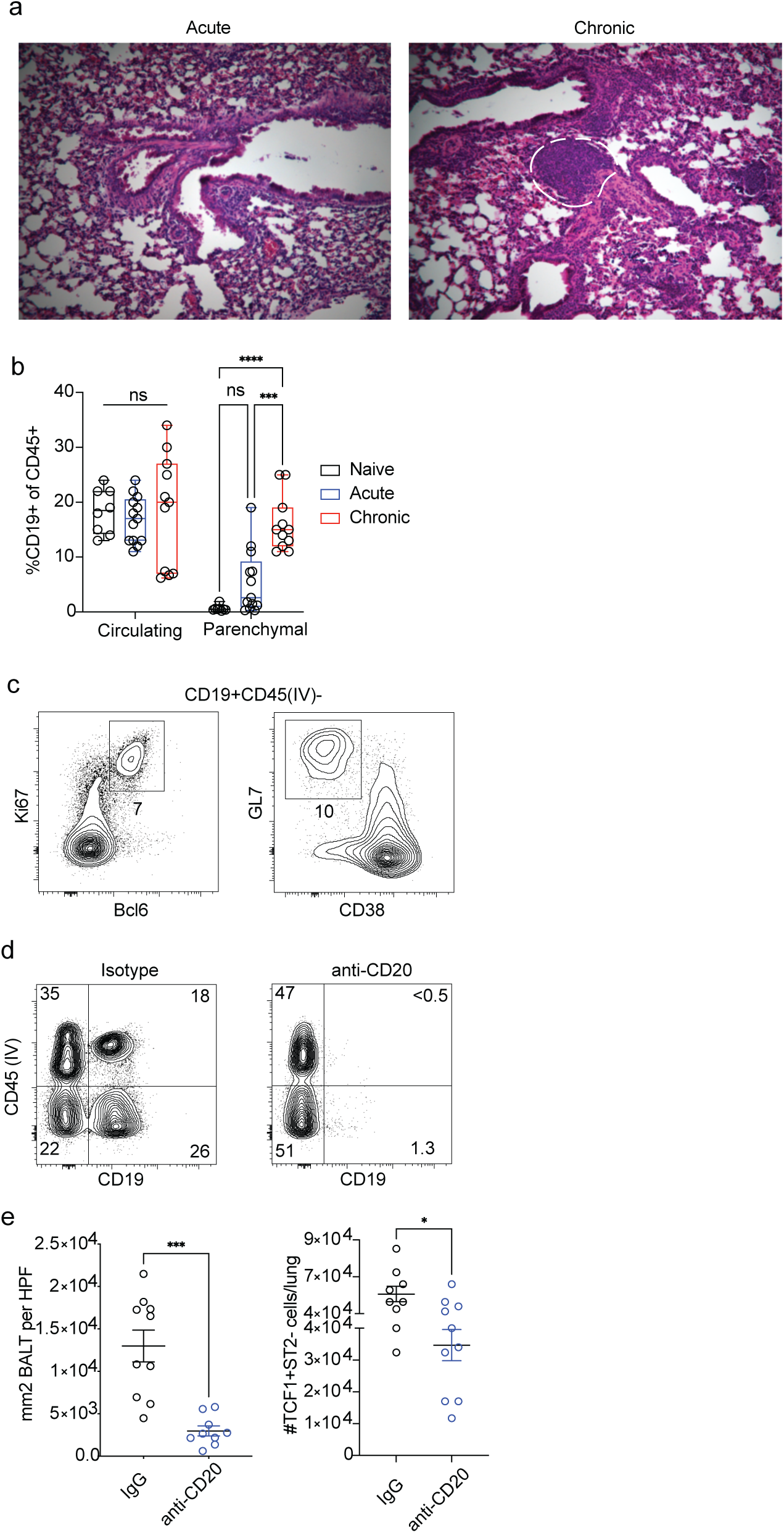
Th2 progenitor cell maintenance is associated with ectopic lymphoid structures and partially dependent on B cells. a) Representative H&E staining of lungs from indicated conditions, with bronchus-associated ectopic lymphoid structure highlighted at the chronic timepoint. b) Frequencies of circulating and lung parenchymal B cells at indicated timepoints. ***p<0.001. ****p<0.0001. n.s. not significant. One-way ANOVA with Holm-Sidak’s correction for multiple comparisons. c) Identification of germinal center B cells by Ki67/Bcl6 intracellular staining and GL7/CD38 surface marker profile at the chronic timepoint of sensitization. FACS plots are representative of 5 biological replicates. d) Frequencies of circulating (CD45IV+CD19+) and parenchymal (CD45IV-CD19+) B cells at chronic timepoint after treatment with isotype control (left) or CD20 depleting antibody (right). FACS plots are representative of 4 biological replicates. e) Left: Quantification of BALT area as square millimeters per high powered field, Right: Quantification of total numbers of Gata3+TCF1+ST2-Th2 progenitors, after treatment with isotype control or CD20 depleting antibody. *p<0.05. ***p<0.001. Unpaired t test.

**Extended Data Figure 4:**
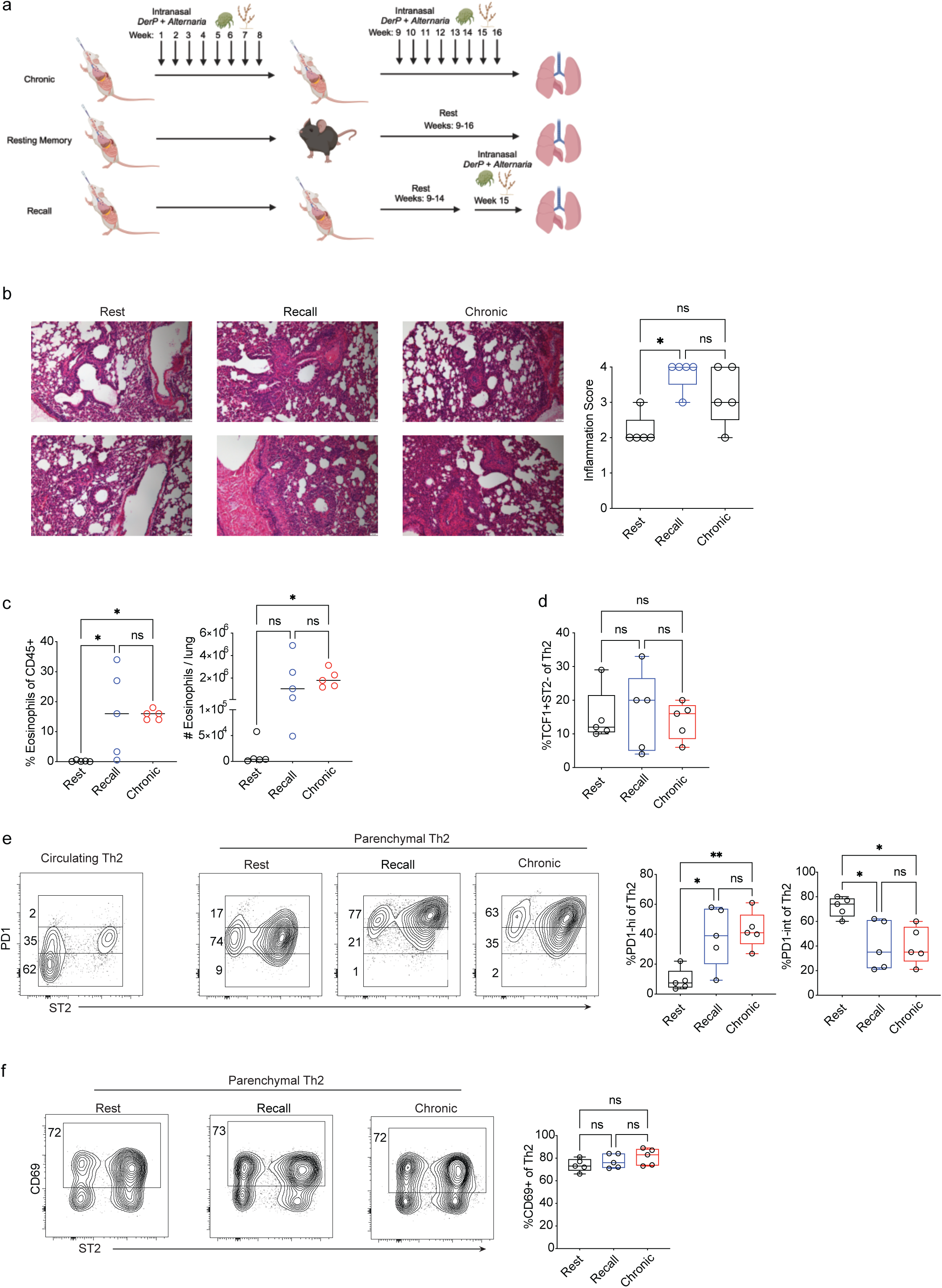
Distinct phenotype of chronic Th2 progenitor cells in comparison with resting and recall tissue resident memory Th2. a) Schematic of chronic, resting memory, and recall models of pulmonary type 2 inflammation b) Left: representative H&E staining of lungs from indicated conditions. Right: quantification of mean inflammation score. *p<0.05, N.s. not significant. One-way ANOVA with Holm-Sidak’s correction for multiple comparisons. c) Quantification of eosinophilic lung infiltrate. Left: % eosinophils of parenchymal (CD45(IV)-) CD45+ immune cells. Right: numbers of eosinophils per lung for indicated conditions. *p<0.05. n.s. not significant. One-way ANOVA with Holm-Sidak’s correction for multiple comparisons. d) %TCF1+ST2-progenitors of Th2 cells e) Representative flow cytometry gating of PD1lo, PD1int, and PD1hi cells in the circulation (left) and lung parenchyma (middle 3 panels) of indicated conditions. Right: quantification of PD1hi and PD1int cell frequencies of total Th2 cells for indicated conditions. *p<0.05, **p<0.01, n.s. not significant. One-way ANOVA with Holm-Sidak’s correction for multiple comparisons. f) Representative flow cytometry gating of CD69 expression by parenchymal Th2 cells in the indicated conditions. n.s. not significant. One-way ANOVA with Holm-Sidak’s correction for multiple comparisons.

**Extended Data Figure 5:**
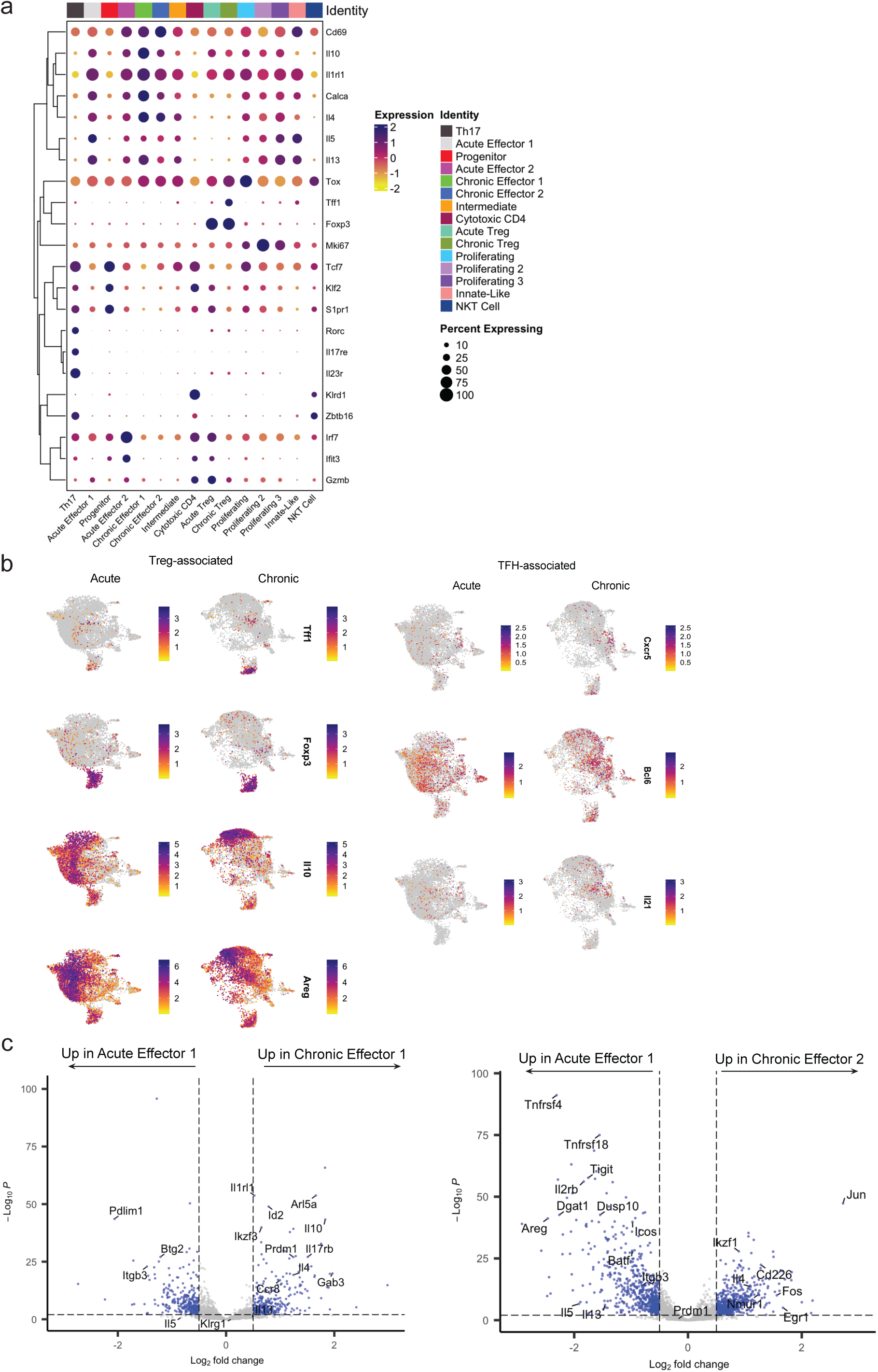
Transcriptomic analysis of CD4+ T cells during acute and chronic Alternaria/DerP sensitization. a) DotPlot of key lineage defining transcripts for each cluster from Figure 3a. b) FeaturePlots of key Treg-associated transcripts (left) and TFH-associated transcripts (right) at acute and chronic timepoints. c) Pseudobulk differential expression analysis of Acute Effector 1 cluster cells compared to Chronic Effector 1 (left) and Chronic Effector 2 (right) by DeSEQ2. Vertical dashed line denotes fold change cutoff of 0.5. Horizontal dashed line denotes significance cutoff of 0.01 by False Discovery Rate (FDR).

**Extended Data Figure 6:**
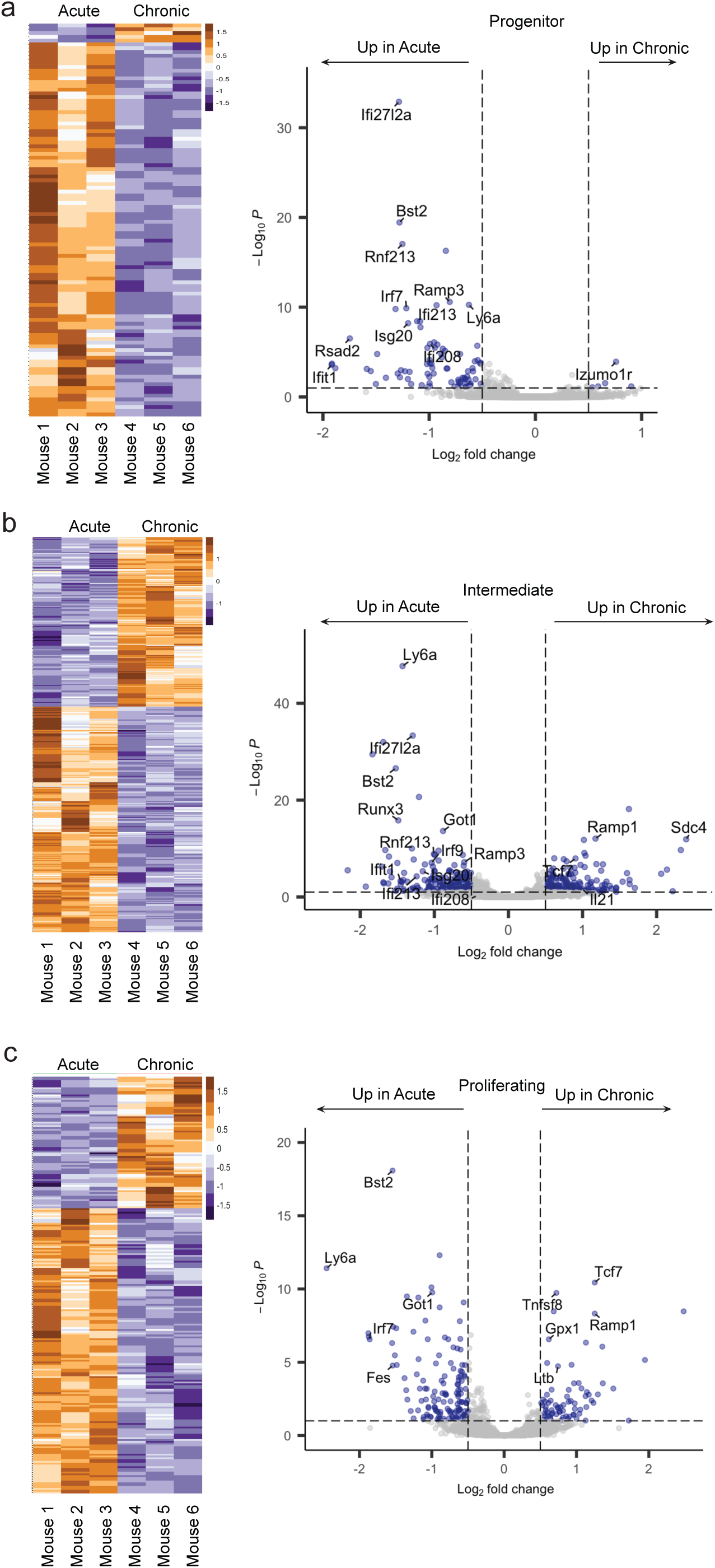
Pseudobulk differential expression analysis of key Th2 clusters during acute and chronic Alternaria/DerP sensitization. Pseudobulk differential expression analysis of acute versus chronic timepoints for a) Progenitor, Intermediate, and c) Proliferating clusters. Vertical dashed line denotes fold change cutoff of 0.5. Horizontal dashed line denotes significance cutoff of 0.01 by False Discovery Rate (FDR).

**Extended Data Figure 7:**
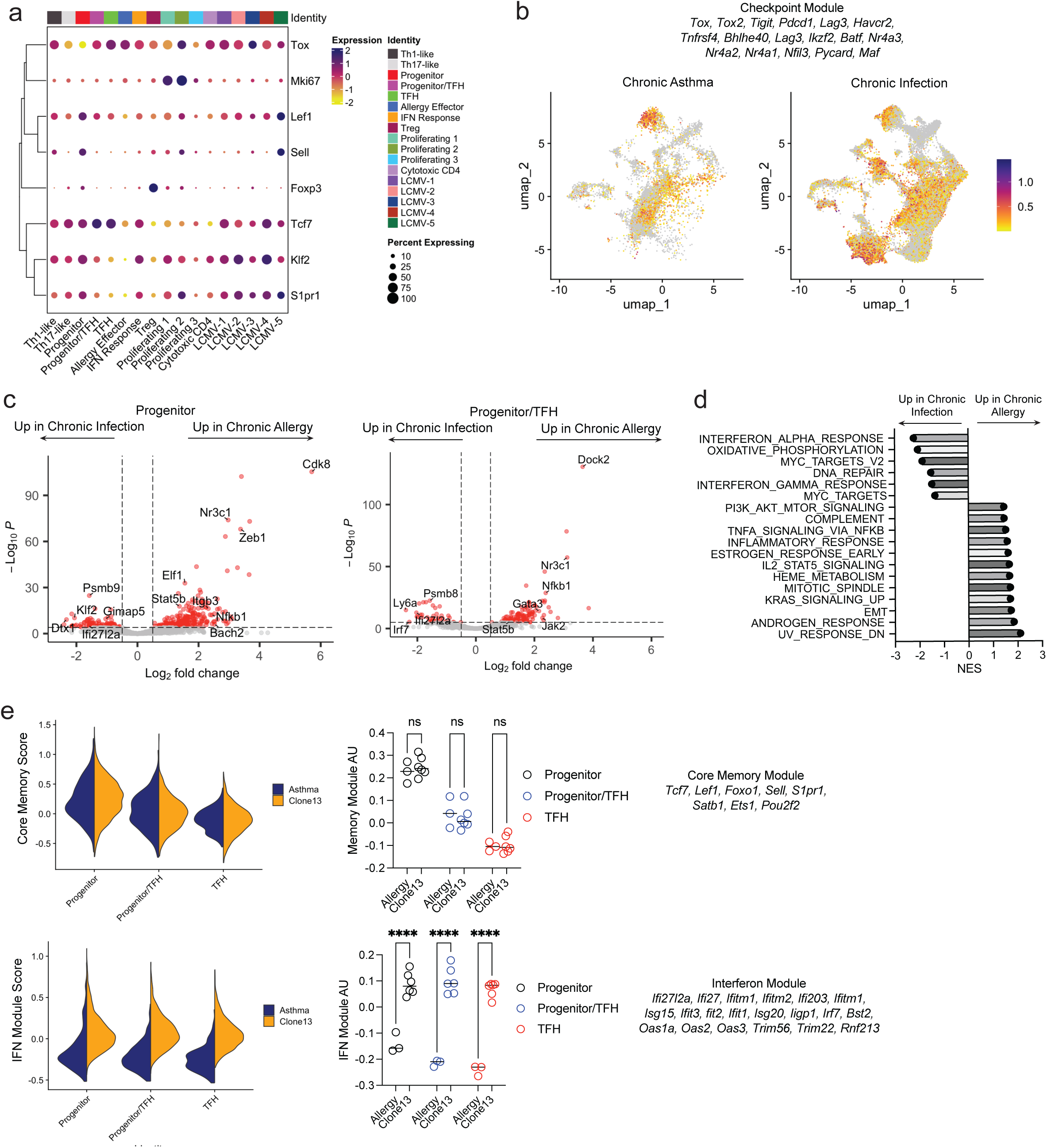
Integrated transcriptomic analysis of CD4+ T cell progenitors in chronic infection and chronic asthma. a) DotPlot of key memory/stemness-associated transcripts for all clusters. b) FeaturePlot of Checkpoint module expression, split by condition (chronic infection vs. chronic asthma). c) Pseudobulk differentiation expression analysis by DeSEQ2 of chronic infection versus chronic asthma for the Progenitor (left) and Progenitor/TFH (right) clusters, displayed as a volcano plot. Vertical dashed line denotes fold change cutoff of 0.5. Horizontal dashed line denotes significance cutoff of 0.01 by False Discovery Rate (FDR). d) Hallmark pathway analysis of differentially expressed transcripts from panel C progenitors with significance p<0.01. e) Upper left: Split violinplots of Core Memory module score for Progenitor, Progenitor/TFH, and TFH clusters, split by condition (chronic infection vs. chronic asthma). Right: quantification of Core Memory module score for indicated clusters. Lower left: Split violinplots of Interferon module score for Progenitor, Progenitor/TFH, and TFH clusters, split by condition (chronic infection vs. chronic asthma). Right: quantification of Interferon module score for indicated clusters. ****p<0.0001. n.s. not significant. One-way ANOVA with Holm-Sidak’s correction for multiple comparisons.

**Extended Data Figure 8:**
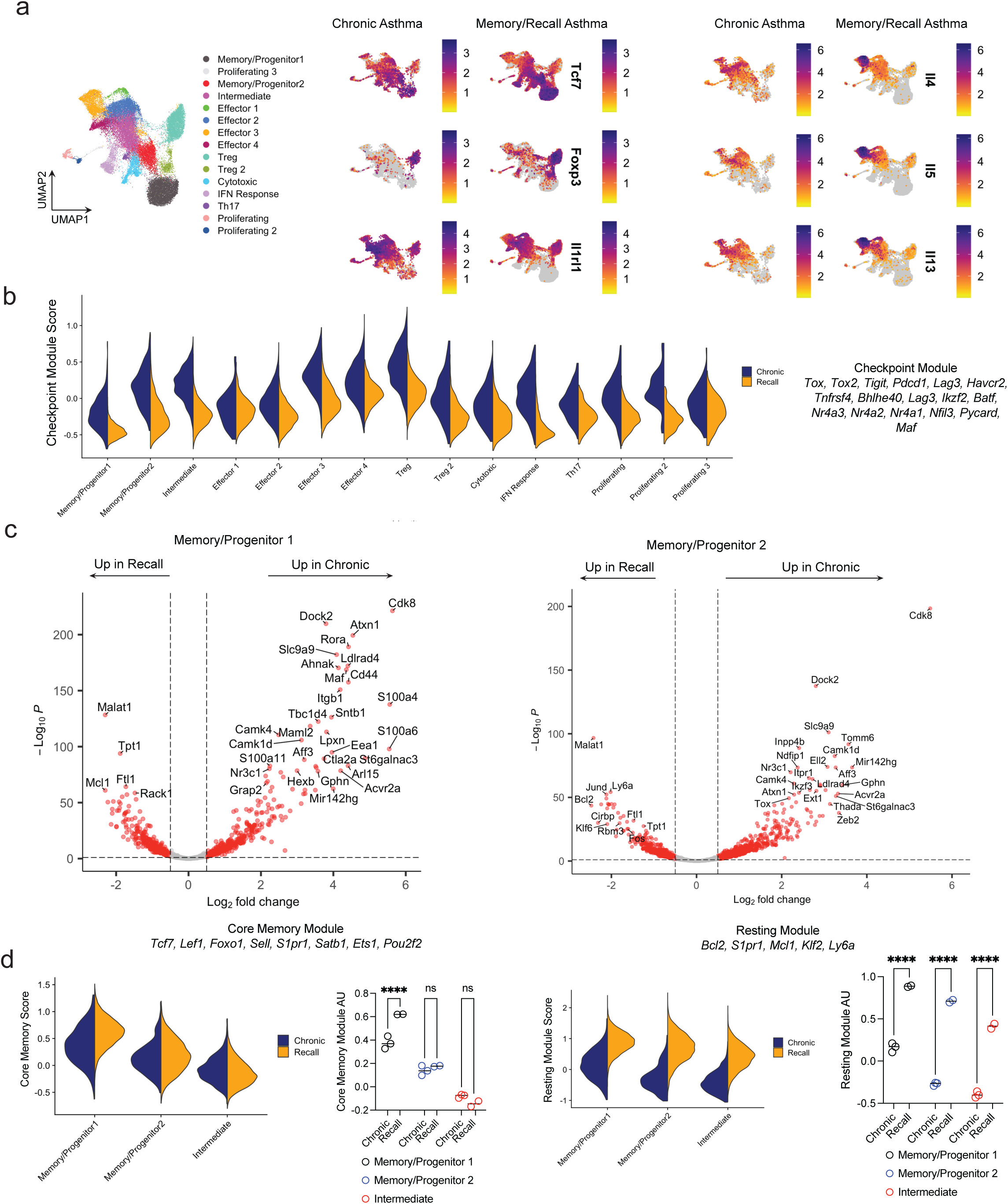
Integrated transcriptomic analysis of Th2 in chronic asthma and resting memory/recall asthma models. a) Left: Integrated UMAP of cells from chronic *Alternaria*/DerP asthma model (Fig 3a) and resting memory/recall *A. Fumigatus* model from Ulrich *et al.*^9^ Right: FeaturePlots of key transcripts associated with Th2 memory and effector cells. b) Split violinplot of Checkpoint module for all clusters. c) Pseudobulk differentiation expression analysis by DeSEQ2 of chronic asthma versus memory/recall for the indicated clusters, displayed as a volcano plot. Vertical dashed line denotes fold change cutoff of 0.5. Horizontal dashed line denotes significance cutoff of 0.01 by False Discovery Rate (FDR). d) Split violinplots of Core Memory module and Resting module, with associated quantification. ****p<0.0001, n.s. not significant. One-way ANOVA with Holm-Sidak’s correction for multiple comparisons.

**Extended Data Figure 9:**
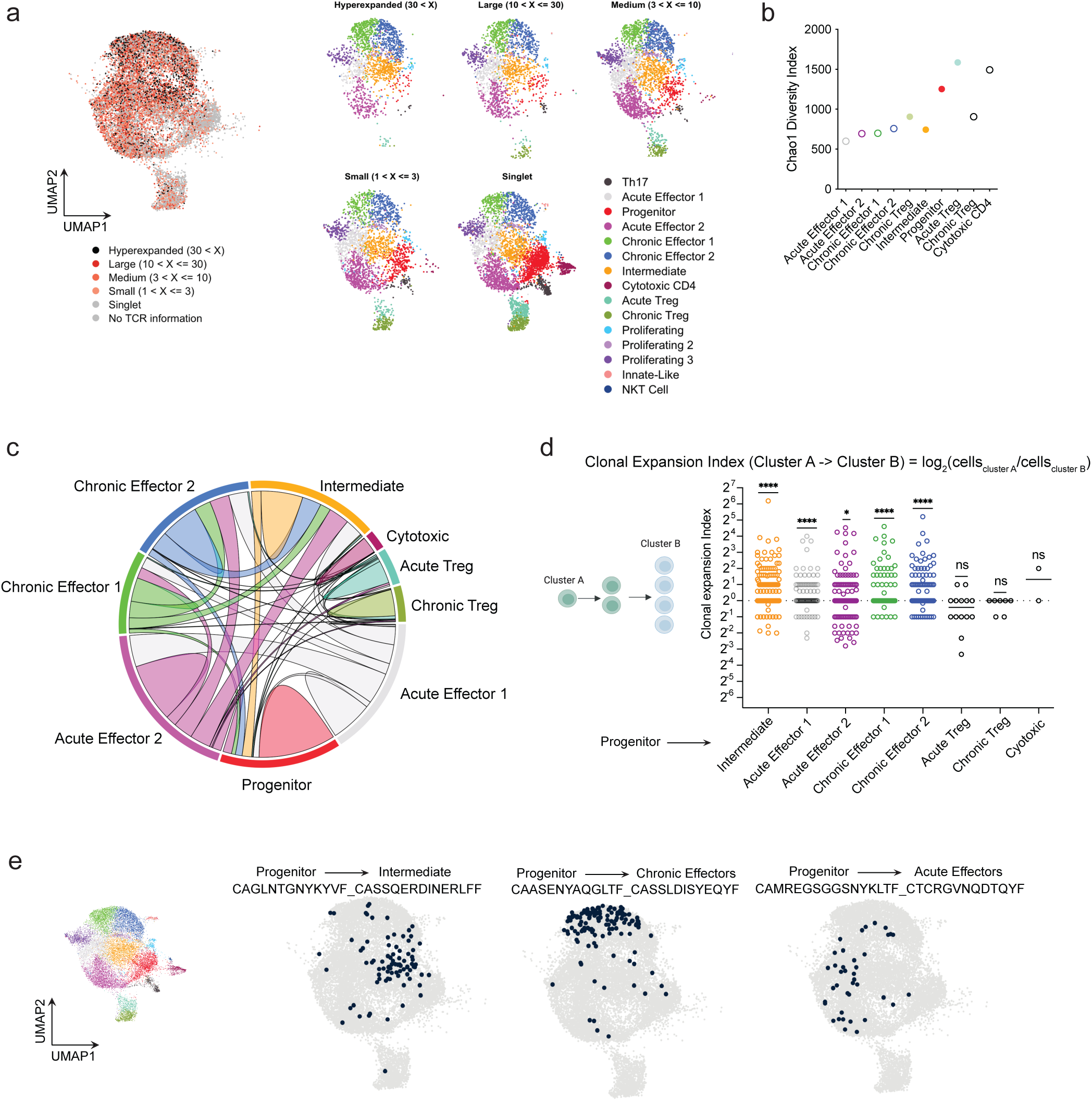
scTCRseq analysis identifies clonal relationships between Th2 subsets. a) Left: UMAP of all cells from acute and chronic timepoints of *Alternaria*/DerP sensitization, colored by degree of clonal expansion. Right: UMAP split by degree of clonal expansion and colored by original cluster colors from Fig 3a. b) Chao clonal diversity index for each ab T cell cluster. c) Circos plot showing clonal overlap between ab T cell clusters. d) Quantification of clonal expansion indices with Progenitor cells as the root node. *p<0.05, ***p<0.0001, n.s. not significant. One sample T test. e) UMAP with individual TCR clones highlighted in black (all other cells light grey), showing clonal expansion between Progenitor and indicated effector cell clusters.

**Extended Data Figure 10:**
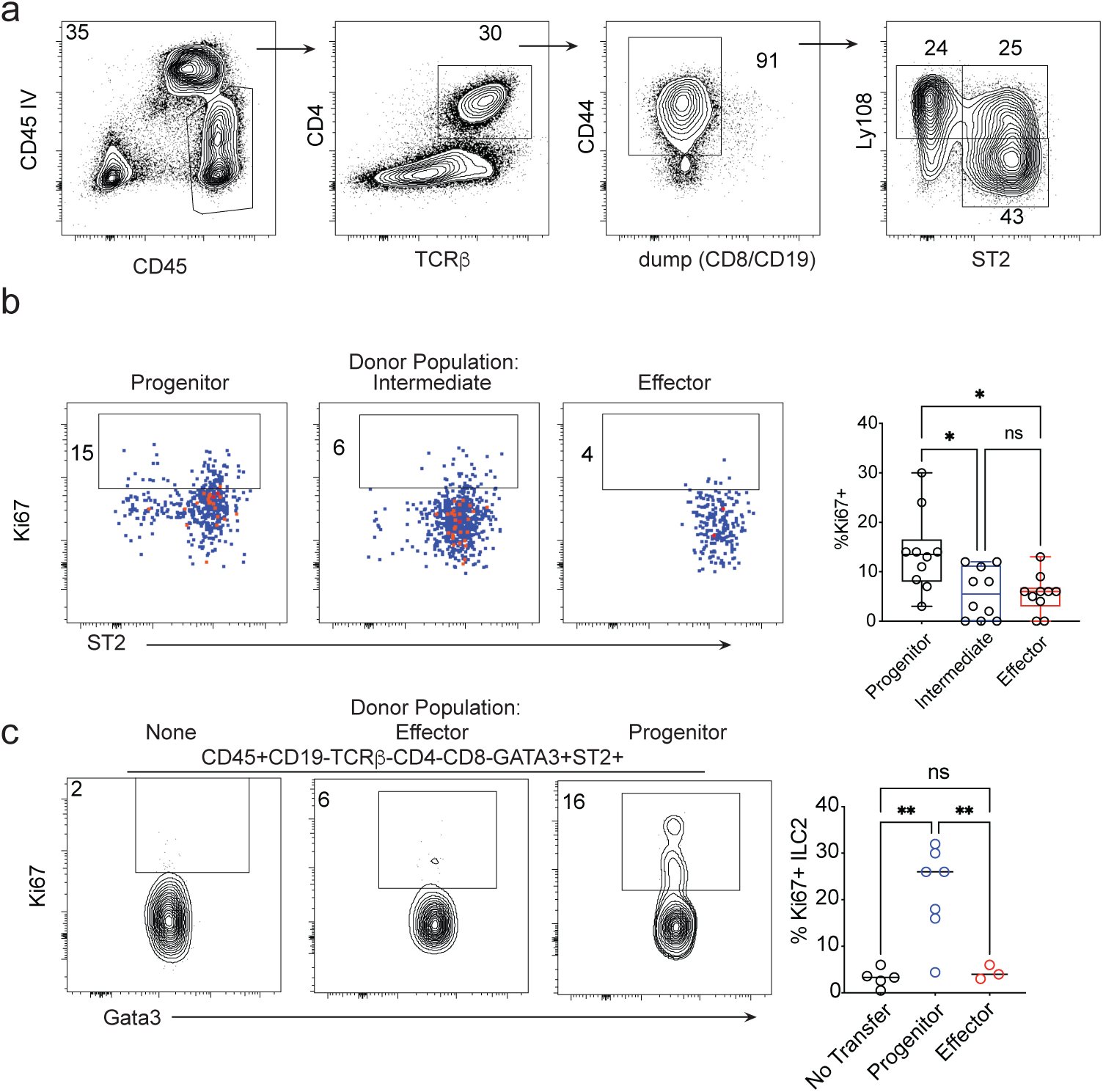
Adoptive transfer of Th2 subsets into TCRbKO recipient mice. a) Representative gating sequence for FACS sorting of Th2 progenitor, intermediate, and effector subsets. b) Left: expression of Ki67 and ST2 by transferred cells of the indicated populations. Right: quantification of Ki67+ Th2 cells. *p<0.05, n.s. not significant. One-way ANOVA with Holm-Sidak’s correction for multiple comparisons. c) Left: quantification of Ki67+ ILC2s after adoptive transfer of the indicated Th2 subsets. Right: *p<0.05, **p<0.01, n.s. not significant. One-way ANOVA with Holm-Sidak’s correction for multiple comparisons.

